# Pan-cancer analysis demonstrates that integrating polygenic risk scores with modifiable risk factors improves risk prediction

**DOI:** 10.1101/2020.01.28.922088

**Authors:** Linda Kachuri, Rebecca E. Graff, Karl Smith-Byrne, Travis J. Meyers, Sara R. Rashkin, Elad Ziv, John S. Witte, Mattias Johansson

## Abstract

Cancer risk is determined by a complex interplay of environmental and heritable factors. Polygenic risk scores (PRS) provide a personalized genetic susceptibility profile that may be leveraged for disease prediction. Using data from the UK Biobank (413,753 individuals; 22,755 incident cancer cases), we quantify the added predictive value of integrating cancer-specific PRS with family history and modifiable risk factors for 16 cancers. We show that incorporating PRS measurably improves prediction accuracy for most cancers, but the magnitude of this improvement varies substantially. We also demonstrate that stratifying on levels of PRS identifies significantly divergent 5-year risk trajectories after accounting for family history and modifiable risk factors. At the population level, the top 20% of the PRS distribution accounts for 4.0% to 30.3% of incident cancer cases, exceeding the impact of many lifestyle-related factors. In summary, this study illustrates the potential for improving cancer risk assessment by integrating genetic risk scores.

## INTRODUCTION

Cancer susceptibility is inherently complex, but it is well-accepted that heritable genetic factors and modifiable exposures contribute to cancer development. While our knowledge of causal modifiable risk factors has gradually evolved over the past decades, genome-wide association studies (GWAS) have rapidly produced a wealth of germline genetic risk variants for different cancers. These studies have shed light on genetic mechanisms of cancer susceptibility, however, the public health impact of GWAS findings has been modest. In response, GWAS results have been leveraged to create polygenic risk scores (PRS) by combining weighted genotypes for risk alleles into a single, integrated measure of an individual’s genetic predisposition to a specific phenotypic profile. Such genetic risk scores are not designed to reflect the complexity of molecular susceptibility mechanisms, but they are highly amenable to phenotypic prediction.

Multiple studies have demonstrated that PRS can generate informative predictions for heritable traits^1, 2^ and diseases^3, 4^, prompting many to advocate for increased integration of genetic risk scores into clinical practice^5, 6^. An important step towards realizing the promise of PRS in precision medicine lies in systematically assessing the added value of genetic information in comparison to conventional risk factors and examining how it affects lifetime risk trajectories^6^. The recent development of large, prospective cohorts with both genome-wide genotyping and deep phenotyping data, such as the UK Biobank^7^, provide an opportunity for integrative analyses of genetic variation and modifiable risk factors. In addition to evaluating PRS predictive performance, these data also provide a unique opportunity to answer etiological questions about the relative contribution of genetic and modifiable risk factors to cancer susceptibility.

In this study we assembled PRS for 16 cancer types, based on previously published GWAS, and applied them to an external population of 413,870 UK Biobank (UKB) participants, with the aim of quantifying the potential of low-penetrance susceptibility variants to improve cancer risk assessment at the population level. First, we assessed the degree to which PRS can improve risk prediction and stratification based on established cancer risk factors, such as family history and modifiable health-related characteristics. Next, we estimated the proportion of incident cancer cases that can be attributed to high genetic susceptibility, captured by the PRS, and compared this to modifiable determinants of cancer. Taken together, our results show that genetic susceptibility captured by the PRS accounts for a substantial proportion of the overall cancer incidence and that incorporating this genetic information improves risk prediction models based on conventional risk factors alone for most cancers.

## RESULTS

### Associations with known risk factors

Characteristics of the UKB study population are presented in Supplementary Table 1. Over the course of the follow-up period a total of 22,755 incident cancers were diagnosed in 413,753 individuals, after excluding participants outside of the age enrollment criteria and those who withdrew consent after enrollment. Established cancer risk factors (listed in Supplementary Table 2) exhibited associations of expected magnitude and direction with each cancer (Supplementary Table 3). Family history of cancer in first-degree relatives, at the corresponding site, conferred a significantly higher risk of prostate (HR=1.84, 95% CI: 1.68-2.00, p=9.1×10^-46^), breast (HR=1.56, 1.44-1.69, p=3.0×10^-29^), lung (HR=1.61, 1.43-1.81, p=7.4×10^-15^), and colorectal (HR=1.26, 1.14-1.40, p=1.2×10^-5^) cancers. Metrics of tobacco use, such as smoking status, intensity, and duration were positively associated with risks of lung, colorectal, bladder, kidney, pancreatic, and oral cavity/oropharyngeal cancers. Weekly alcohol intake was associated with higher risks of breast (HR per 70 grams = 1.04, p=2.3×10^-5^), colorectal (HR=1.04, p=5.9×10^-9^), and oral cavity/pharyngeal (HR=1.05, p=3.0×10^-10^) cancers. Adiposity was associated with cancer risk at multiple sites, including endometrium (BMI: HR per 1-unit = 1.09, 1.08-1.10, p=1.6×10^-49^), colon/rectum (waist-to-hip ratio: HR per 10% increase = 1.17, 1.11-1.24, p=2.2×10^-8^), and kidney (BMI: HR=1.04, 1.02-1.05, p=1.7×10^-6^). Particulate matter (PM_2.5_) was associated with lung cancer risk^8^ (PM_2.5_: HR per 1 micro-g/m^3^ = 1.10, 1.05-1.15, p=1.9×10^-5^) in the model that included smoking status and intensity.

All PRS associations with the target cancer reached at least nominal statistical significance (Figure 1; Supplementary Table 4). We considered three PRS approaches (see Methods for details): standard weights corresponding to reported risk allele effect sizes (PRS_β_); unweighted sum of risk alleles (PRS_unw_); inverse variance weights that incorporate the standard error of the risk effect size (PRS_IV_). The latter approach resulted in stronger or equivalent (HR ± 0.01) associations for most cancers, except Non-Hodgkin lymphoma (NHL). Compared to standard PRS_β_, substantial differences were observed for prostate (PRS_IV_: HR=1.77, *P*=4.3×10^-366^ vs. PRS_β_: HR=1.39, *P*=2.0×10^-105^), colon/rectum (PRS_IV_: HR=1.48, *P*=1.8×10^-94^ vs. PRS_β_: HR=1.32, *P*=5.5×10^-50^), leukemia (PRS_IV_: HR=1.70, *P*=6.3×10^-23^ vs. PRS_β_: HR=1.45, *P*=8.0×10^-13^), and thyroid (PRS_IV_: HR=1.75, *P*=1.9×10^-15^ vs. PRS_β_: HR=1.57, *P*=5.7×10^-10^). All subsequent analyses use PRS_IV_ since this approach appears to improve PRS performance by appropriately downweighing the contribution of variants with less precisely estimated effects.

**Figure 1:**
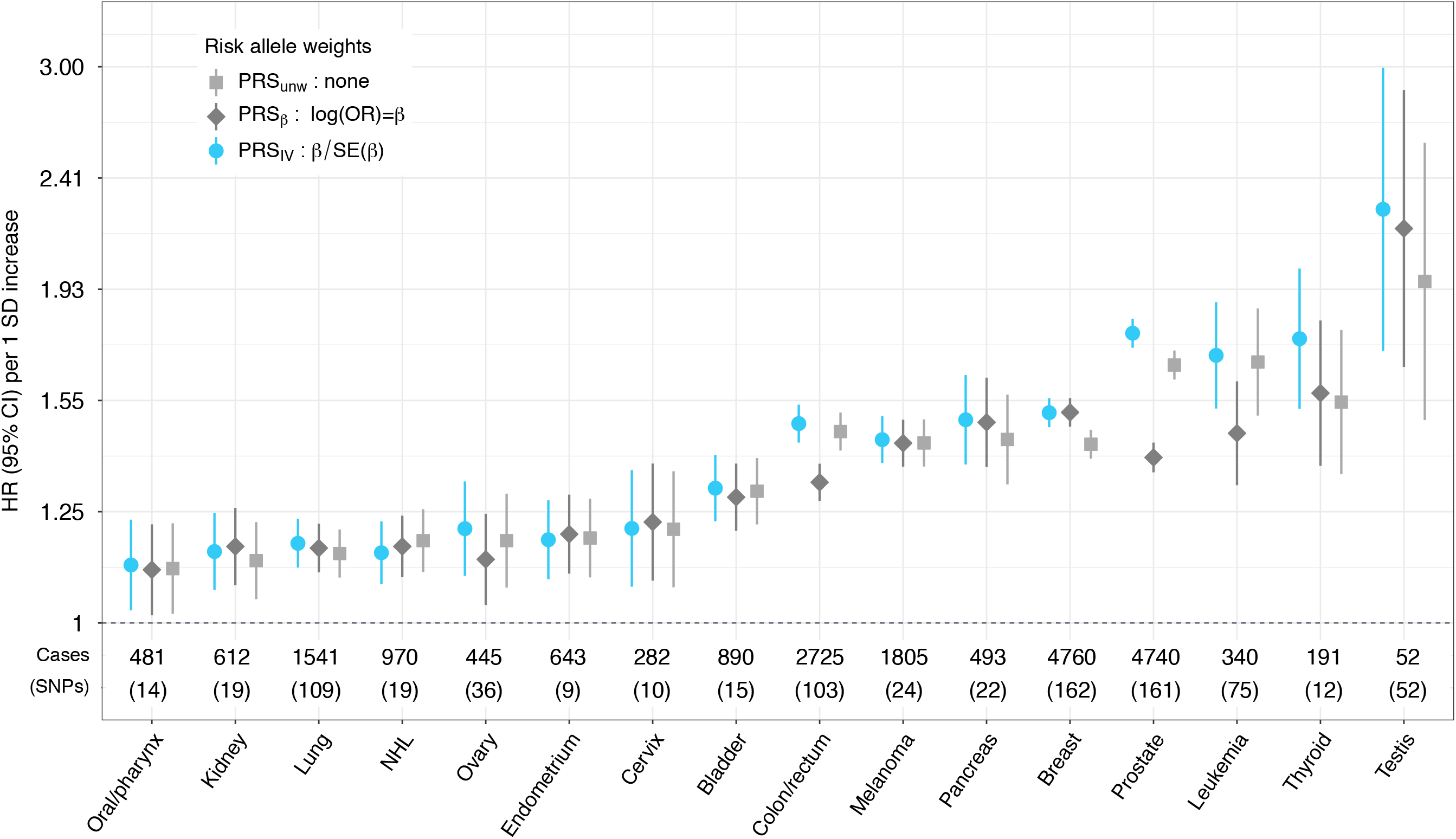
Hazard ratios (HR) per one standard deviation (SD) increase in the standardized polygenic risk score (PRS). HR were estimated using cause-specific Cox proportional hazards models, accounting for mortality as a competing risk. A comparison of three weighting approaches for combining individual risk variants in the PRS is presented: standard weights based on per-allele log odds ratios (PRS_β_), unweighted sum of risk alleles (PRS_unw_), and inverse variance (IV) weights (PRS_IV_).

### Improvement in risk prediction

The predictive performance of each risk model was evaluated based on its ability to accurately estimate risk (calibration) and distinguish cancer cases from cancer-free individuals (discrimination). All cancer-specific risk models were well-calibrated (Goodness of fit p>0.05; Supplementary Figure 1). Model discrimination was assessed by Harrell’s C-index, estimated as a weighted mean between 1 and 5 years of follow-up time. For completeness, we also report the AUC at 5 years of follow-up time^9^. Proportionality violations (p<0.05) were detected for age in the breast cancer model and PRS_IV_ for cervical cancer. For breast cancer this was resolved by incorporating an interaction term with follow-up time. As a sensitivity analysis for cervical cancer we modelled a time-varying PRS effect (Supplementary Figure 2).

The C-index reached 0.60 with age and/or sex, for all cancers except for breast and thyroid (Supplementary Table 5). For cancers with available information on family history of cancer at the same site (prostate, breast, colon/rectum, and lung), incorporating this had a modest impact on the C-index (ΔC<0.01). In fact, replacing family history with the PRS resulted in an improvement in discrimination for prostate (C=0.763, ΔC=0.047), breast (C=0.620, ΔC=0.061), and colorectal (C=0.708, ΔC=0.029), but not lung (C=0.711, ΔC=-0.002) cancers.

Next, we assessed the change in the C-index (ΔC) after incorporating the PRS into prediction models with all available risk factors for each cancer (Figure 2; Supplementary Table 5). The resulting improvement in prediction performance was variable. The largest increases in the C-index were observed for cancer sites with few available predictors, such as testes (C_PRS_=0.766, ΔC=0.138), thyroid (C_PRS_=0.692, ΔC=0.099), prostate (C_PRS_=0.768, ΔC=0.051) and lymphocytic leukemia (C_PRS_=0.756, ΔC=0.061). Incorporating the PRS also improved prediction accuracy for melanoma (C_PRS_=0.664, ΔC=0.042), breast (C_PRS_=0.635, ΔC=0.063), and colorectal (C_PRS_=0.716, ΔC=0.030) cancers, which have multiple environmental risk factors. The highest overall C-index was observed for lung (C_PRS_=0.849) and bladder (C_PRS_=0.814) cancers, which was primarily attributed to non-genetic predictors (C without PRS: lung = 0.846; bladder = 0.808). However, it is worthwhile noting that despite having a large ΔC, the precision of the C-index estimates was low some for rarer cancers, such as testicular (n=52) and thyroid (n=191), as well as cancers with genetic risk scores based on relatively few variants. Changes in the AUC at 5 years of follow-up were of similar magnitude (Supplementary Table 5).

**Figure 2:**
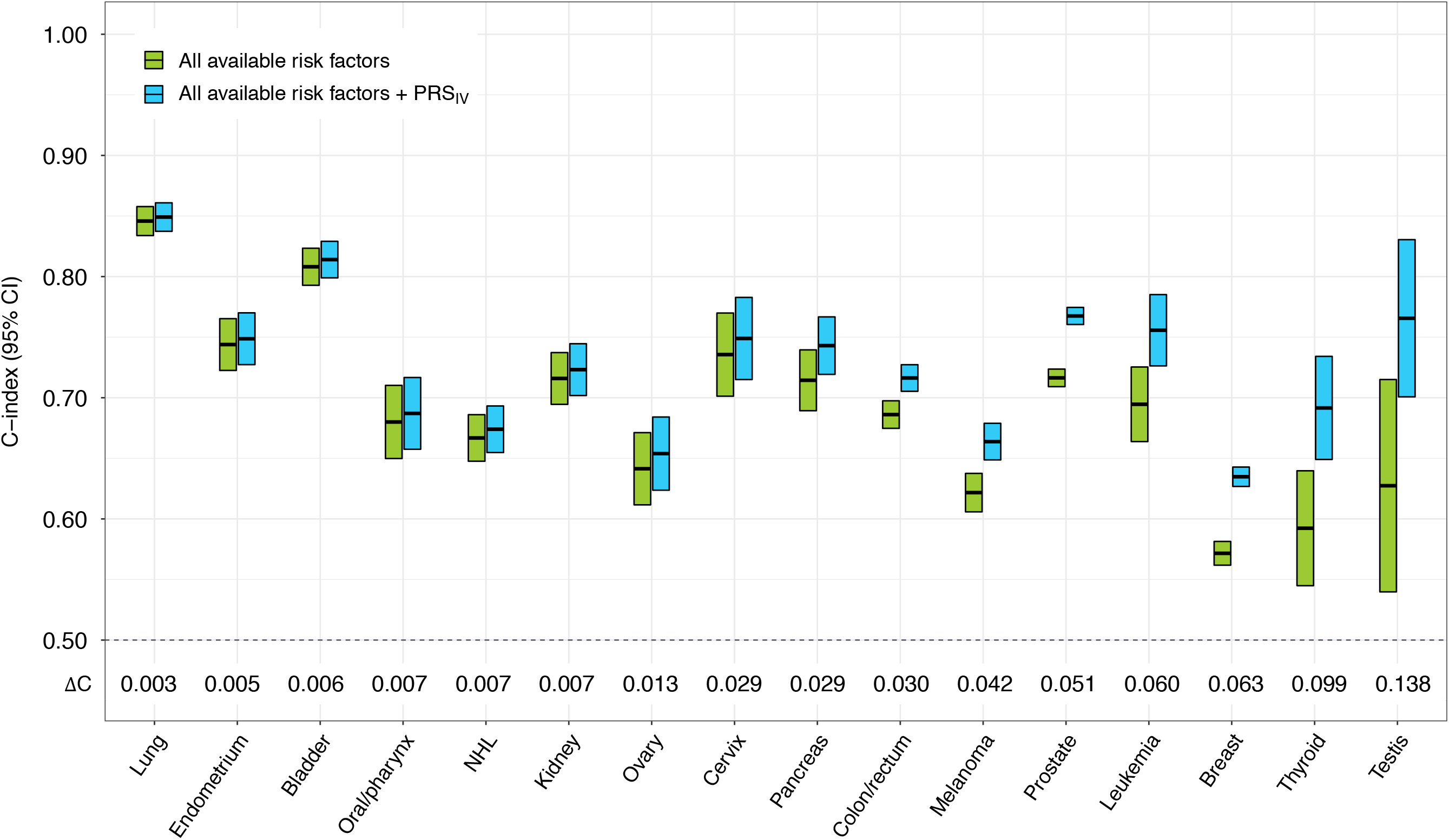
Assessment of model discrimination based on Harrell’s C index between 1 and 5 years of follow-up time. Comparisons are conducted between the most comprehensive risk factor model for each cancer, including all available lifestyle-related risk factors and family history (if applicable), and a nested model that also includes the standardized polygenic risk score (PRS_IV_) for that cancer and the top 15 genetic ancestry principal components.

As a complementary metric of model performance, Royston’s R^2^ was calculated to quantify the variation in the time-to-event outcome captured by each risk model^10^. Across all 16 sites, the median change in R^2^ (ΔR^2^) was 0.066. Large improvements, defined as ΔR^2^ >0.10, were observed for cancers of the breast (R^2^_PRS_=0.146; ΔR^2^=0.103), pancreas (R^2^_PRS_=0.439; ΔR^2^=0.103), leukemia (R^2^_PRS_=0.415; ΔR^2^=0.160), prostate (R^2^_PRS_=0.510; DR^2^ =0.161), thyroid (R^2^_PRS_=0.310; DR^2^ =0.230), and testis (R^2^_PRS_=0.605; DR^2^ =0.421). These results parallel the trend in improvement observed based on C-index and AUC.

For 15 out of 16 cancers, incorporating the PRS resulted in significant improvement in reclassification, as indicated by positive percentile-based net reclassification index (NRI)^11^ values with 95% bootstrapped confidence intervals excluding 0 (Supplementary Table 6). The overall NRI was primarily driven by the event NRI (NRI_e_), which is the increase in the proportion of cancer cases reclassified to a higher risk group. Positive NRI_e_ values >0.25 were observed for prostate, thyroid, breast, testicular, leukemia, melanoma, and colorectal cancers. The largest reclassification improvement in non-event NRI (NRI_ne_) observed for the lung PRS (NRI_ne_=0.015) and breast PRS (NRI_ne_=0.014). Four cancers (testes, leukemia, kidney, oral cavity/pharynx) had significantly negative NRI_ne_ values indicating that adding the PRS decreased classification accuracy in cancer-free individuals.

### Refinement of risk stratification

The ability of the PRS to refine risk estimates was assessed by examining 5-year absolute risk trajectories as a function of age, across strata defined by percentiles of PRS (high risk ≥80%, average: >20% to <80%, low risk: ≤20%) and family history of cancer (Figure 3; exact p-values in Supplementary Table 7). Significantly diverging risk trajectories, overall and at age 60, were observed for prostate (*P*≤4.5×10^-25^), breast (*P*≤4.6×10^-32^), colorectal (*P*≤2.0×10^-21^), and lung cancers (*P*≤0.031). For all cancers except lung, risk stratification was primarily driven by PRS. For instance, 60-year-old men with a high PRS but no family history of prostate cancer had a higher mean 5-year disease risk (4.74%) compared to men with a positive family history and an average PRS (3.66%). For lung cancer, on the other hand, participants with a positive family history had higher average 5-year risks, even with a low PRS (0.54%), compared to those without (high PRS: 0.46%; low PRS: 0.29%). There was evidence of interaction between the PRS and family history of cancer for prostate *(P* = 9.0×10^-128^), breast *(P* = 1.2×10^-98^), colorectal *(P* = 8.7×10^-14^) cancers (Supplementary Table 8). For lung cancer the interaction with family history was limited to the high PRS group (*P* = 5.9×10^-3^).

**Figure 3:**
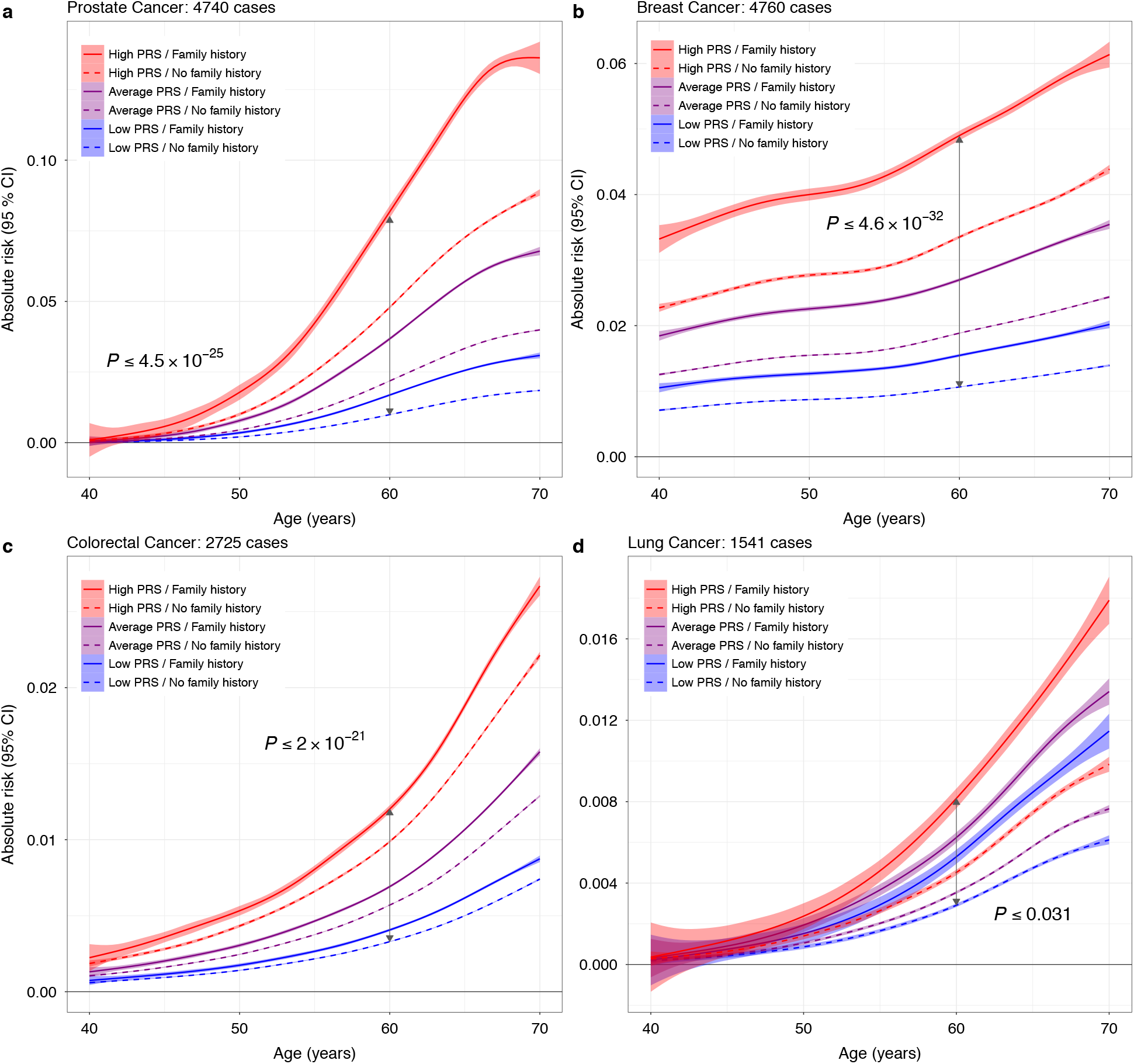
Predicted 5-year absolute risk trajectories across strata defined by family history and percentiles of the polygenic risk score (PRS) distribution. Family history was based on self-reported illnesses in first-degree relatives for **(a)** prostate, **(b)** breast, **(c)** colorectal, and **(d)** lung cancers. Low PRS corresponds to ≤20^th^ percentile, average PRS is defined as >20^th^ to <80^th^ percentile, and high PRS includes individuals in the ≥80^th^ percentile of the normalized genetic risk score distribution. P-values for differences in mean absolute risk in each stratum at age 60 are based on t-tests (two-sided).

We also compared 5-year risk projections across strata of PRS and modifiable risk factors. Effects of multiple risk factors were combined into a single score by generating summary linear predictors for each cancer (see Methods for details). For several common cancers, individuals with a high PRS were predicted to have an overall risk above the median, and this increased risk was observed even when high PRS individuals also had modifiable risk factor scores that were below the median modifiable risk factor score (Figure 4; Supplementary Figure 3). PRS achieved significant risk stratification for breast cancer (premenopausal: *P*≤7.9×10^-20^; post-menopausal: *P*≤1.7×10^-40^), colorectal cancer (*P*≤1.8×10^-42^), and melanoma (*P*≤3.5×10^-139^) (Figure 4; exact p-values in Supplementary Table 7). The same pattern of stratification was observed for NHL, leukemia, pancreatic, thyroid, and testicular cancers (Supplementary Figures 3-4). For other phenotypes, lifestyle-related risk factors had a stronger overall influence on risk trajectories than PRS (Figure 5; exact p-values in Supplementary Table 7). However, the stratifying by levels of PRS still resulted in significantly diverging risk projections for several cancers (lung: *P*≤1.1×10^-13^; oral cavity/pharynx: *P*≤1.2×10^-12^; kidney: *P*≤1.7×10^-52^). For bladder cancer, the risk trajectories for high PRS/reduced modifiable risk and low PRS/high modifiable risk were overlapping (*P*=0.99).

**Figure 4:**
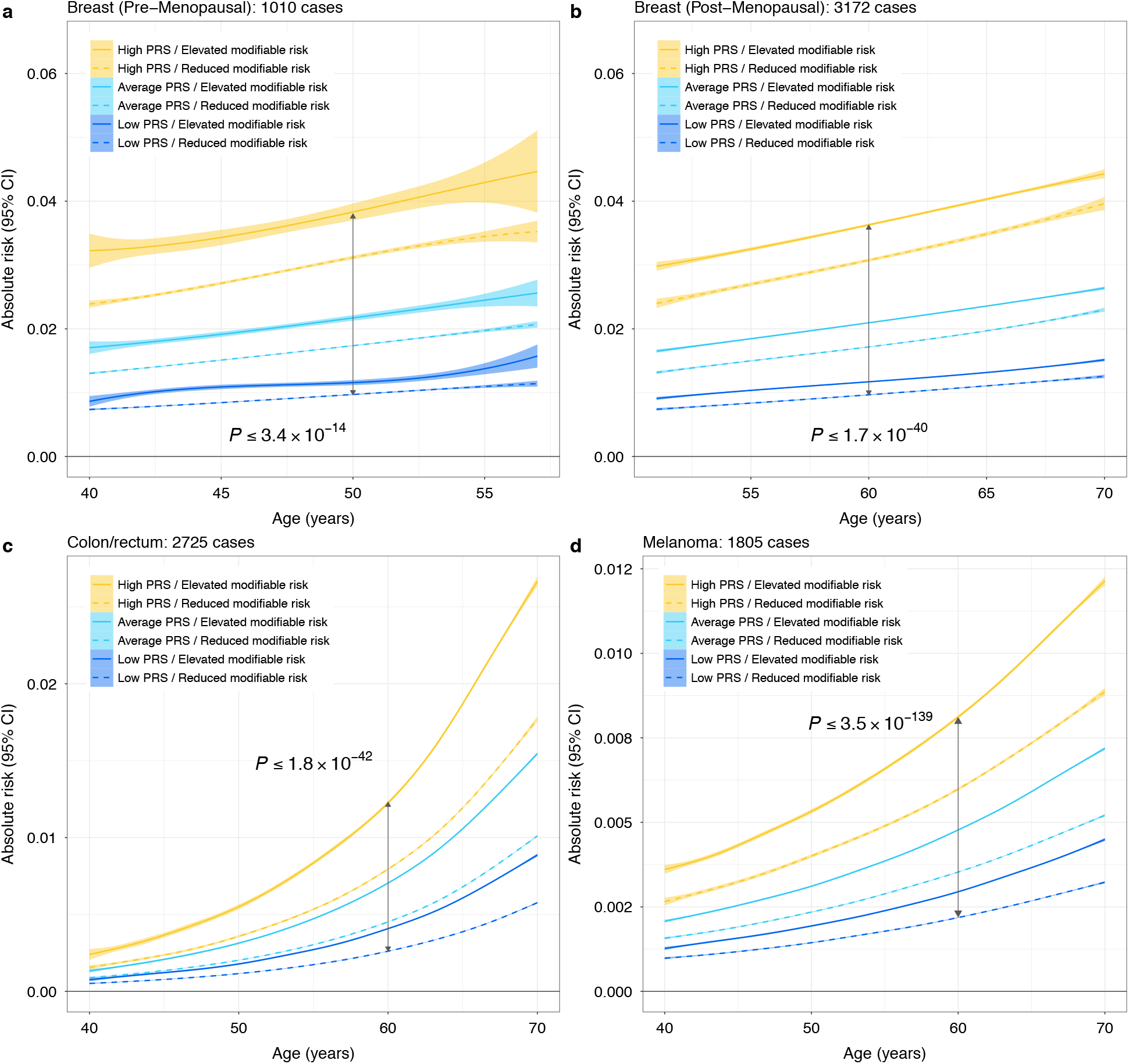
Predicted 5-year absolute risk trajectories across strata defined by modifiable risk factors and percentiles of the polygenic risk score (PRS) distribution. Low PRS corresponds to ≤20^th^ percentile, average PRS is defined as >20^th^ to <80^th^ percentile, and high PRS includes individuals in the ≥80^th^ percentile of the normalized genetic risk score distribution. Individuals below the median of the modifiable risk factor distribution were classified as having reduced risk, whereas those above the median had elevated risk. P-values for differences in mean absolute risk in each stratum at age 50 for **(a)** pre-menopausal breast cancer and at age 60 for **(b)** post-menopausal breast cancer, **(c)** colon/rectal cancer, and **(d)** melanoma are based on t-tests (two-sided).

**Figure 5:**
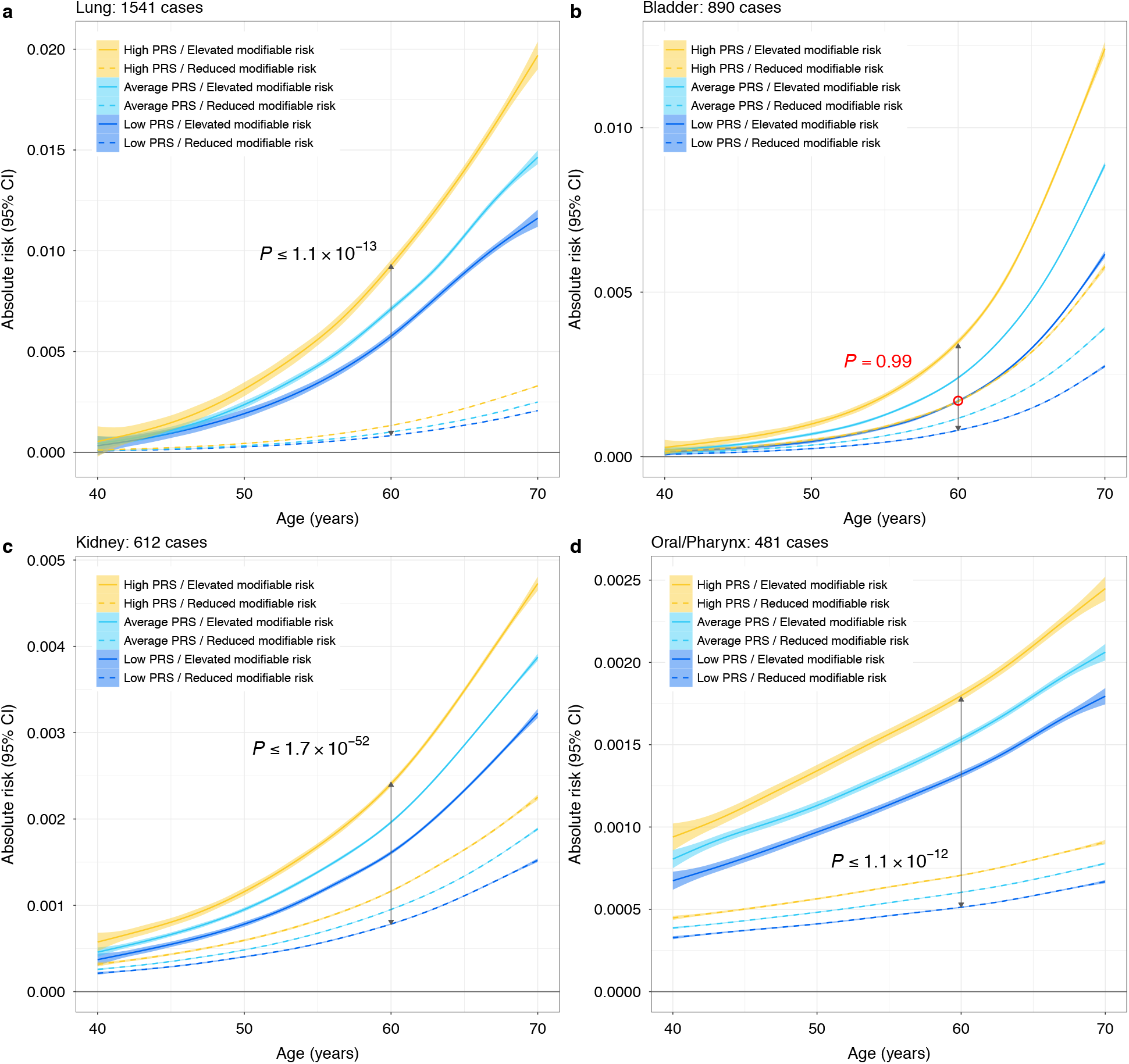
Predicted 5-year absolute risk trajectories across strata defined by modifiable risk factors and percentiles of the polygenic risk score (PRS) distribution. Low PRS corresponds to ≤20^th^ percentile, average PRS is defined as >20^th^ to <80^th^ percentile, and high PRS includes individuals in the ≥80^th^ percentile of the normalized genetic risk score distribution. Individuals below the median of the modifiable risk factor distribution were classified as having reduced risk, whereas those above the median had elevated risk. P-values for differences in mean absolute risk in each stratum at age 60 for **(a)** lung, **(b)** bladder, **(c)** kidney, and **(d)** oral/pharynx cancers are based on t-tests (two-sided).

There was evidence of larger than additive risk differences, at age 60 between elevated modifiable risk factor profiles and all ordinal PRS categories for melanoma (*P*=3.3×10^-122^), breast cancer (postmenopausal: *P*=6.9×10^-24^; pre-menopausal: *P*=4.9×10^-7^, colorectal (*P*=1.3×10^-208^), lung (*P*=1.1×10^-37^), bladder (*P*=1.5×10^-50^), kidney (*P*=5.5×10^-29^), and oral cavity/pharynx cancers (*P*=5.2×10^-11^) (Supplementary Table 8).

### Quantifying population-level impact

Population attributable fractions (PAF) were used to summarize the relative contribution of genetic susceptibility and modifiable risk factors to cancer risk at the population level. In order to allow comparisons between PAF estimates, the PRS and modifiable risk score distributions were both dichotomized at ≥80^th^ percentile. All risk factors nominally contributed (*P*<0.05) to cancer incidence (Figure 6; Supplementary Table 9), with the exception of the PRS for oral cavity/pharynx cancer (*P*=0.78) and PM2.5 for lung cancer in never smokers (*P*=0.44).

**Figure 6:**
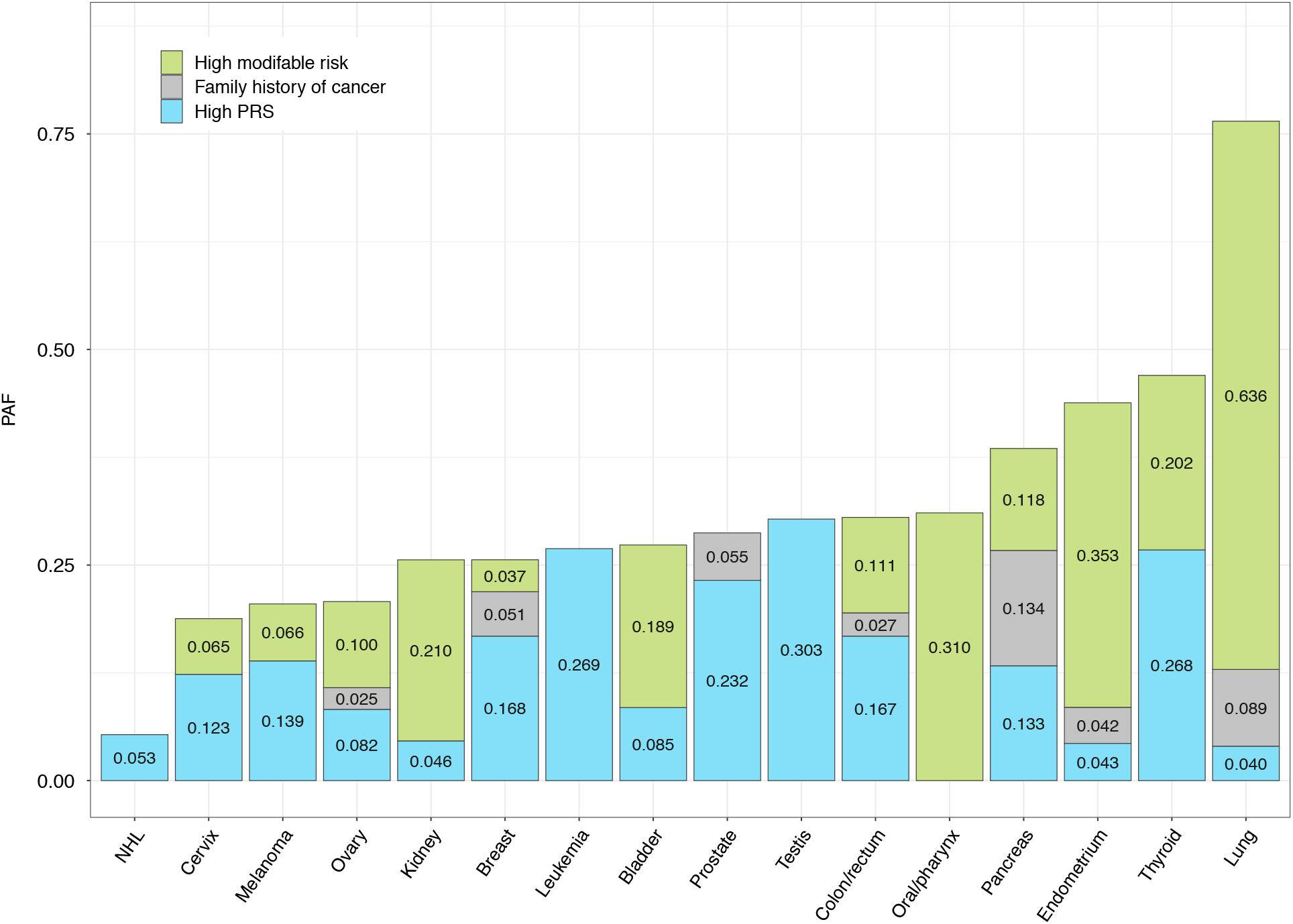
Population attributable fractions (PAF) estimated at 5 years of follow-up time for the top 20% (≥80^th^ percentile) of the modifiable risk factor and polygenic risk score (PRS) distributions, respectively, and family history of cancer at the relevant site. PAF estimates were derived from Cox proportional hazard regression models that were adjusted for age at enrollment, sex, family history of cancer (if available), genotyping array, and the top 15 genetic ancestry principal components.

PAF for high genetic risk exceeded the contribution of modifiable exposures for several cancers, such as thyroid (PAF_PRS_=0.268, *P*=1.7×10^-9^), prostate (PAF_PRS_=0.232, *P*=5.5×10^-158^), colon/rectum (PAF_PRS_=0.167, *P*=9.2×10^-50^), breast (PAF_PRS_=0.168, *P*=4.9×10^-87^), and melanoma (PAF_PRS_=0.139, *P*=1.3×10^-23^). For testicular cancer (PAF_PRS_=0.303, *P*=4.5×10^-4^), leukemia (PAF_PRS_=0.269, *P*=4.5×10^-4^), lung cancer in never smokers (PAF_PRS_=0.077, *P*=0.045), and NHL (PAF_PRS_=0.053, *P*=1.9×10^-3^), PRS was the only significant risk factor other than demographic factors. Cancers for which modifiable risk factors had a substantially larger impact on disease burden than PRS included oral cavity/pharynx (PAF_mod_=0.310 vs. PAF_PRS_=0.006), lung (AF_mod_=0.636 vs. PAF_PRS_=0.040), endometrium (PAF_mod_=0.353 vs. PAF_PRS_=0.043), kidney (PAF_mod_=0.210 vs. PAF_PRS_=0.046), and bladder cancers (PAF_mod_=0.189 vs. PAF_PRS_=0.085). For other sites, such as pancreas (PAF_mod_=0.118 vs. PAF_PRS_=0.133) and ovary (PAF_mod_=0.100 vs. PAF_PRS_=0.082), the contribution of PRS and modifiable risk factors were more balanced.

## DISCUSSION

Cancer is a multifactorial disease with a complex web of etiological factors, from macro-level determinants, such as health policy, to individual-level characteristics, such as health-related behaviors and heritable genetic profiles. Heritable and modifiable risk factors act in concert to influence cancer development, but their relative contributions to disease risk are rarely compared directly in the same population. In this study we provide insight into the potential utility of PRS for cancer risk prediction and provide insight into the relative of contribution of genetic and modifiable risk factors to cancer incidence at population level.

Our first major finding is that cancer-specific PRS comprised of lead GWAS variants improve risk prediction for all 16 cancers examined. However, the magnitude of the resulting improvement in prediction varies substantially between sites. In evaluating the added predictive value of the PRS it is important to keep in mind that achieving the same incremental increase in the C-index/AUC is more difficult when the baseline model already performs well^12^. This was applicable to most cancers, where age and/or sex alone achieved non-trivial risk discrimination (C-index/AUC>0.60). Expanding the set of predictors to include modifiable risk factors further improved discrimination, as previously shown^13^. By adding the PRS to the most comprehensive risk factor models facilitated by our data, we adopted a conservative approach for quantifying its added predictive value, which provides an informative benchmark for future efforts seeking to incorporate genetic predisposition in cancer risk assessment.

Cancer sites for which the PRS resulted in the largest gains in prediction performance included prostate, testicular, and thyroid cancers, as well as leukemia, and melanoma. This is consistent with high heritability estimates reported for these cancers in twin studies^14^ and our analyses in the UK Biobank^15^. Modelling the PRS in addition to established risk factors yielded very modest improvements in risk discrimination for cancers of the lung, endometrium, bladder, oral cavity/pharynx, and kidney. These cancers have strong environmental risk factors, such as smoking, alcohol consumption, obesity, and HPV infection, some of which were captured in our analysis. Limited predictive ability for cervical and endometrial cancers may also be due to a low number of variants included in the PRS (9 and 10, respectively). The association of the lung cancer PRS with cigarettes per day^16^ may have diminished its apparent predictive value when added to a model with smoking status and intensity, which already achieved an AUC>0.80 making difficult to elicit further improvement. Furthermore, PRS may be particularly relevant for assessing lung cancer risk in never smokers, since other risk factors have a limited impact in this population.

Few pan-cancer PRS studies have been conducted in prospective cohorts and none have considered the breadth of modifiable risk factors that we evaluated. Shi et al.^17^ tested 11 cancer PRS in cases from The Cancer Genome Atlas and controls from the Electronic Medical Records and Genomics Network. This analysis was limited by fewer risk variants in each PRS, as well as potential for bias due to selection of cases and controls from different populations. A phenome-wide analysis in the Michigan Genomics Initiative cohort by Fritsche et al.^18^ examined PRS for 12 cancers and reported similar associations for the target phenotype. However, risk stratification was not formally evaluated. Considering cancer-specific studies, the PRS presented here achieved superior prediction performance for some cancers^19, 20, 21, 22^, but not others^23, 24^. For pancreatic cancer^25^ and melanoma^26^, our results are consistent with previous analyses using PRS of similar composition. Generally, comparison of prediction performance is complicated by differences in PRS composition, population characteristics, and inclusion of other predictors. Outside the cancer literature, our conclusions align with a recent study of ischemic stroke, which demonstrated that the PRS is similarly or more predictive than multiple established risk factors, including family history^27^.

Our second major finding advances the idea of using germline genetic information to refine individual risk estimates. We show that incorporating PRS improves risk stratification provided by conventional risk factors alone, as illustrated by significantly diverging 5-year risk projections within strata based on family history or modifiable risk factors. For certain cancers, including some with strong environmental risk factors, such as melanoma, breast, colorectal, and pancreatic cancers, PRS was the primary determinant of risk stratification. For others, such as lung and bladder cancers, modifiable risk factors had a stronger impact on 5-year risk trajectories. A consistent finding for all cancers was that individuals in the top 20% of the PRS distribution with an unfavorable modifiable risk factor profile had the highest level of risk, with evidence that the effects of PRS and modifiable risk factors may be synergistic. Similar risk stratification results based on genetic and modifiable risk factors have also been reported for coronary disease^28^ and Alzheimer’s^29^. Taken together, our results suggest that PRS can provide more accurate risk estimates for individuals with wide variation in cancer predisposition based on lifestyle-related risk factors. Furthermore, as a risk factor that is present and stable throughout the life course, PRS may be useful for motivating targeted prevention efforts in high-risk individuals before they accumulate a high burden of modifiable risk factors.

In addition to evaluating predictive performance and risk stratification, our work demonstrates the relevance of common genetic risk variants at the population level. High genetic risk (PRS≥80^th^ percentile) explained between 4.0% and 30.3% of incident cancer cases, and for many phenotypes this exceeded PAF estimates for modifiable risk factors or family history. The contribution of genetic variation to disease risk is typically conveyed by heritability, which is an informative metric, although not easily translated into a measure of disease burden useful in a public health context. Recent work on cancer PAF in the UK^30^ and a series of publications from the ComPARe initiative in Canada^31, 32^ examined wide range of modifiable risk factors. Despite providing useful data, these studies overlook the contribution of genetic susceptibility. Our work addresses these limitations by providing a more complete perspective on the determinants of cancer and potential impact of future prevention policies.

In evaluating the contributions of our study, several limitations should be acknowledged. First, we did not account for the impact of workplace exposures and socio-economic determinants of health, thereby underestimating the role of non-genetic risk factors. We also lacked data on several known carcinogens, such as ionizing radiation, and clinical biomarkers, such as prostate-specific antigen, thus limiting the extent to which our results inform risk classification for certain cancers. Information on family history was also not available for all cancer types. Second, since the UK Biobank cohort is unrepresentative of the general UK population due to low participation and resulting healthy volunteer bias^33^, we may have underestimated PAFs for modifiable risk factors. Finally, the models presented here are calibrated to the UKB population and we urge caution in extrapolating prediction performance and absolute risk projections to other populations. Since our analytic sample is restricted to individuals of predominantly European ancestry, this limits the applicability of our findings to diverse populations.

This work has several important strengths. Our study provides a comprehensive description of the joint and relative influence of genetic and modifiable risk factors in a population-based cohort with uniform phenotyping and extensive data on a range of relevant cancer risk factors. We established risk models based on the current knowledge of genetic and modifiable risk factors and report a series of metrics that comprehensively characterize different dimensions of PRS predictive performance in an independent population. With the exception of limited overlap with one colorectal cancer GWAS (see Methods), all of our risk models were developed based on previously published associations from studies that did not include the UK Biobank. While our results are promising, we anticipate that the PRS performance reported here may be enhanced by adopting less stringent p-value thresholding, optimizing subtype-specific weights, and implementing more sophisticated PRS models that incorporate linkage disequilibrium structure, functional annotations, or SNP interactions. Some of these strategies are already being successfully implemented^4, 23^. We also provide insight into PRS modelling by showing that accounting for the variance in risk allele effect sizes improves PRS performance. This approach may be particularly advantageous for PRS derived from multiple sources rather than a single GWAS. Throughout this study we consider a relatively lenient definition of high genetic risk, corresponding to the top 20% of the PRS distribution. Exploring other cut-points will be informative, however, our results are valuable for demonstrating that the utility of PRS for stratification is not limited to the most extreme ends of the genetic susceptibility spectrum. This threshold is also compelling from a population-health perspective, as it allows us to quantify the proportion of cases attributed to a risk factor with a 20% prevalence.

Genetic risk scores have the potential to become a powerful tool for precision health, but only if the resulting information can be understood and acted on appropriately. One important consideration is the accuracy and stability of PRS-based risk classifications, especially at clinically actionable risk thresholds that exist for certain cancers. For instance, there are established screening programs for breast and colorectal cancers, and increasing evidence supporting the effectiveness of low-dose computed tomography for lung cancer screening^34, 35^. For these cancers PRS could be used to adjust the optimal age for screening initiation and/or intensity. However, to justify this, studies are needed to demonstrate the benefit of using PRS to supplement conventional screening criteria. Such trials are already underway for breast cancer, where genetic risk scores are being incorporated to personalize risk-based screening^36^. For other cancers, such as prostate, screening remains controversial and PRS may prove useful in identifying a subset of high-risk individuals who may benefit the most from screening.

Another area where PRS may prove useful is for prioritizing individuals for targeted health and lifestyle-related interventions. In support of this, our study demonstrates that those with the highest levels of genetic risk, based on the PRS, may also experience larger decreases in risk from shifting to a healthier lifestyle. However, there is also accumulating evidence that simply reporting genetic risk information to individuals does not induce behavior change that could lead to meaningful reductions in risk^37^. Therefore, progress in our ability to construct and apply PRS to identify high-risk individuals must be also accompanied by the development of effective behavioral interventions that can be implemented in response to high disease risk, in addition to early detection and screening protocols.

Ultimately, the impact of PRS on clinical decision-making should be carefully evaluated in randomized trials prior to deployment in healthcare settings. By demonstrating cancer-specific improvements in risk prediction, as well as the substantial proportion of cancer incidence that is captured by known genetic susceptibility variants, we provide evidence that contextualizes the potential for using genetic information to improve cancer outcomes.

## METHODS

### Study Population

The UK Biobank (UKB) is a population-based prospective cohort of individuals aged 40 to 69 years, enrolled between 2006 and 2010. All participants completed extensive questionnaires, in-person physical assessments, and provided blood samples for DNA extraction and genotyping^7^. Health-related outcomes were ascertained via individual record linkage to national cancer and mortality registries and hospital inpatient encounters^7^. Individuals with at least one recorded incident diagnosis of a borderline, in situ, or malignant primary cancer were defined as cases. Cancer diagnoses coded by International Classification of Diseases (ICD)-9 or ICD-10 codes were converted into ICD-O-3 codes using the SEER site recode paradigm in order to classify cancers by organ site.

Participants were genotyped on the UKB Affymetrix Axiom array (89%) or the UK BiLEVE array (11%)^7^. Genotype imputation was performed using the Haplotype Reference Consortium as the main reference panel, supplemented with the UK10K and 1000 Genomes phase 3 reference panels^7^. Genetic ancestry principal components (PCs) were computed using fastPCA^38^ based on a set of 407,219 unrelated samples and 147,604 genetic markers^7^. All analyses were restricted to self-reported European ancestry individuals with concordant self-reported and genetically inferred sex. To further minimize potential for population stratification, we excluded individuals with values for either of the first two ancestry PCs outside of five standard deviations of the population mean. Based on a subset of genotyped autosomal variants with minor allele frequency (MAF)≥0.01 and genotype call rate ≥97%, we excluded samples with call rates <97% and/or heterozygosity more than five standard deviations from the mean of the population. With the same subset of SNPs, we used KING^38^ to estimate relatedness among the samples. We excluded one individual from each pair of first-degree relatives, preferentially retaining individuals to maximize the number of cancer cases remaining, resulting in a total of 413,870 UKB participants.

### Polygenic Risk Scores

In order to derive polygenic risk scores (PRS) for each of the 16 cancers, we extracted previously associated variants by searching the National Human Genome Research Institute (NHGRI)-European Bioinformatics Institute (EBI) Catalog of published GWAS. For every eligible GWAS, both the original primary manuscript and supplemental materials were reviewed. Additional relevant studies were identified by examining the reference section of each article and via PubMed searches of other studies in which each article had been cited. We abstracted all autosomal variants with minor allele frequency MAF≥ 0.01 and *P*<5×10^-8^ identified in populations of at least 70% European ancestry and published by June 2018, with the exception of one colorectal cancer GWAS^39^ (published in December 2018). Studies used to identify cancer risk variants and obtain corresponding effect sizes for the PRS were conducted in populations other than the UK Biobank. One exception is the colorectal cancer study by Huyghe et al.^39^, which included 5356 cases and 21,407 controls from the UK Biobank in the GWAS meta-analysis, comprising 9% of cases and 21% of all participants.

Details of the PRS development approach, including a comprehensive list of source studies, is described by Graff et al^16^. For inclusion in the PRS we preferentially selected independent SNPs (LD *r*^2^<0.3) with the highest imputation score and we excluded SNPs with allele mismatches or MAF differences >0.10 relative to the 1000 Genomes reference population, and palindromic SNPs with MAF≥0.45. For associations reported in more than one study of the same ancestry and phenotype, we selected the one with the most information (i.e., which reported the risk allele and effect estimate) with the smallest p-value. For breast cancer, we looked up 187 candidate PRS variants in publicly meta-analysis summary statistics from the Breast Cancer Association Consortium (BCAC) GWAS, as reported in Michalidou et al.^40^. We retained SNPs with *P*<5×10^-8^ in the BCAC meta-analysis (n=162) and constructed a standard PRS using risk allele weights from these summary statistics.

Three approaches for combining risk variants in the PRS were considered. First, we used standard PRS weights, corresponding to the log odds ratio (β) for each risk allele:

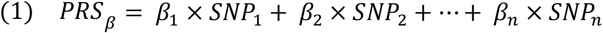

We compared this to an unweighted score corresponding to the sum of the risk alleles, which is equivalent to assigning all variants an equal weight of 1:

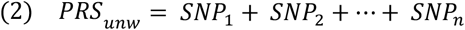

Lastly, we applied inverse variance (IV) weights that incorporated the standard error (SE) of the SNP log(OR) to account for uncertainty in risk allele effect sizes and downweigh the contribution of variants with less precisely estimated associations (weights provided in **Supplementary Data 1**):

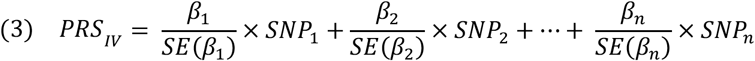

Each PRS was standardized with the entire cohort to have a mean of 0 and standard deviation (SD) of 1.

### Risk Model Development and Evaluation

Cancer-specific prediction models consisting of four classes of risk factors were developed: i) demographic factors (age and sex); ii) family history of cancer in first-degree relatives; iii) modifiable risk factors; and iv) genetic susceptibility, represented by the PRS. Family history of cancer was derived based on self-reported illnesses in non-adopted first-degree relatives, which only listed cancers of the prostate, breast, bowel, or lung. In addition to these four cancer sites, family history of breast cancer was included as a predictor for ovarian cancer^41, 42^. Models for pancreatic cancer included a composite variable for family history of cancer at any of these four sites^43, 44^. Selection of modifiable risk factors was informed by literature review and reports, such as the European Code Against Cancer^45^, IARC Monographs, and evaluations from the World Cancer Research Fund International, with an emphasis on risk factors that are likely to have a causal role. Final models included established environmental and lifestyle-related characteristics that were collected for the entire UK Biobank cohort (**Supplementary Table 1**).

Cause-specific Cox proportional-hazard models were used to estimate the hazard ratios (HR) and corresponding 95% confidence intervals (CI) for genetic and lifestyle factors associated with each incident cancer. Death from any cause, other than cancer site-specific mortality, was treated as a competing event. Information on primary and contributing causes of death was used to identify cancer site-specific mortality. Follow-up time was calculated from the date of enrollment to the date of cancer diagnosis, date of death, or end of follow-up (January 1,2015). For each cancer, individuals with a past or prevalent cancer diagnosis at that same site were excluded from the analysis, while individuals diagnosed with cancers at other sites were retained in the population. All models including the PRS were also adjusted for genotyping array and the first 15 genetic ancestry PCs. For the PRS, HR estimates correspond to 1 SD increase in the standardized genetic score.

The predictive performance of each risk model was evaluated based on its ability to accurately estimate risk (calibration) and distinguish cancer cases from cancer-free individuals (discrimination). Calibration was assessed with a Hosmer-Lemeshow goodness-of-fit statistic modified for time-to-event outcomes^46^, and by plotting the expected event status against the observed event probability^47^ across risk deciles (or quantiles to ensure a minimum of 5 cases per group). Violation of the proportionality of hazards assumption was assessed by examining the association between standardized Schoenfeld residuals and time.

We evaluated nested models starting with the most minimal set of predictors, such as demographic factors, followed by models including family history of cancer and modifiable risk factors, and finally models incorporating the PRS. Risk discrimination was assessed based on Harrell’s C-index, calculated as a weighted average between 1 and 5 years of follow-up time, and Area Under the Curve (AUC) at 5 years. We also report pseudo *R*^2^ coefficients based on Royston’s measure of explained variation for survival models^10^. Percentile-based net reclassification improvement (NRI) index^11^ was used to quantify improvements in reclassification. NRI summarizes the proportion of appropriate directional changes in predicted risks. Any upward movement in risk categories for cases indicates improved classification, and any downward movement implies worse reclassification. The opposite is expected for non-cases:

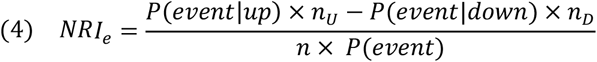

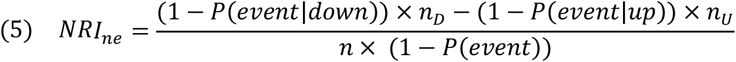

where *n_U_* is the number of individuals up-classified and *n_D_* is the number down-classified. Overall NRI is the sum of the NRI in cases and non-cases: *NRI* = *NRI_e_* + *NRI_ne_*. Bootstrapped confidence intervals were obtained based on 1000 replicates.

### Assessment of Risk Stratification

For each individual, we estimated the 5-year absolute risk of being diagnosed with a specific cancer using the formula of Benichou & Gail^48^, as implemented by Ozenne et al^49^. Absolute risk trajectories were examined as a function of age across strata defined by genetic and modifiable risk profiles, as well as family history. Individuals in the top 20% of the PRS distribution (PRS≥80^th^ percentile) for a given cancer were classified has having high genetic risk, those in the bottom 20% (PRS≤20^th^ percentile) were classified as low risk, and the middle category (>20^th^ to <80^th^ percentile) classified as average genetic risk.

Modifiable risk factors were summarized by generating summary linear predictors (predicted log-hazard ratios) based on risk factors in **Supplementary Table 1**, excluding age, sex, and family history. Individuals above the median of this risk score distribution were considered to have an unfavorable modifiable risk profile. Risk trajectories in each stratum were visualized by fitting linear models with smoothing splines to individual risk estimates as a function of age. Differences in mean risk at age 60 were tested using a two-sample t-test. We also tested for interaction between the 3-level ordinal PRS variable and the modifiable risk score (dichotomized at the median) in a linear model with the predicted absolute risk as the outcome.

The relative contribution of genetic and modifiable cancer risk factors at the population level was quantified with population attributable fractions (PAF) using the method of Sjölander & Vansteedlandt^50, 51^ based on the counterfactual framework. To obtain comparable AF estimates, thresholds for high genetic risk and high burden of modifiable risk factors corresponded to the top 20% (≥80^th^ percentile) of each risk score distribution.

## DATA AVAILABILITY

The UK Biobank in an open access resource, available at https://www.ukbiobank.ac.uk/researchers/. This research was conducted with approved access to UK Biobank data under application number 14105. All the other data supporting the findings of this study are available within the article and its supplementary information files and from the corresponding author upon reasonable request. A reporting summary for this article is available as a Supplementary Information file.

## URLs

R package RiskRegression: https://cran.r-project.org/web/packages/riskRegression/index.html

Polygenic risk scores available from: https://www.pgscatalog.org/publication/PGP000050/

## ACKNOWLEDGEMENTS

This research was supported by funding from the National Institutes of Health (US NCI R25T CA112355 and R01 CA201358; PI: Witte) and Cancer Research UK (C18281/A19169).

## AUTHOR CONTRIBUTIONS

Study conception: L.K., K.S.B., M.J.; Development of analytic strategy: L.K., K.S.B., J.S.W., M.J.; Polygenic risk score data acquisition: R.E.G., T.J.M.; Statistical analysis: L.K., R.E.G.; UK Biobank genotype and sample quality control: S.R.R., R.E.G; Project coordination: M.J., J.S.W.; L.K drafted the manuscript with input from K.S.B, M.J., J.S.W, E.Z.; All authors contributed to the interpretation of the results and provided critical feedback on the manuscript.

## References

1. Khera AV, et al. Polygenic Prediction of Weight and Obesity Trajectories from Birth to Adulthood. Cell 177, 587–596 e589 (2019).

2. Yengo L, et al. Meta-analysis of genome-wide association studies for height and body mass index in approximately 700000 individuals of European ancestry. Hum Mol Genet 27, 3641–3649 (2018).

3. Inouye M, et al. Genomic Risk Prediction of Coronary Artery Disease in 480,000 Adults: Implications for Primary Prevention. J Am Coll Cardiol 72, 1883–1893 (2018).

4. Khera AV, et al. Genome-wide polygenic scores for common diseases identify individuals with risk equivalent to monogenic mutations. Nat Genet 50, 1219–1224 (2018).

5. Torkamani A, Wineinger NE, Topol EJ. The personal and clinical utility of polygenic risk scores. Nat Rev Genet 19, 581–590 (2018).

6. Lambert SA, Abraham G, Inouye M. Towards clinical utility of polygenic risk scores. Hum Mol Genet, (2019).

7. Bycroft C, et al. The UK Biobank resource with deep phenotyping and genomic data. Nature 562, 203–209 (2018).

8. Raaschou-Nielsen O, et al. Air pollution and lung cancer incidence in 17 European cohorts: prospective analyses from the European Study of Cohorts for Air Pollution Effects (ESCAPE). Lancet Oncol 14, 813–822 (2013).

9. Heagerty PJ, Zheng Y. Survival model predictive accuracy and ROC curves. Biometrics 61, 92–105 (2005).

10. Royston P. Explained variation for survival models. The Stata Journal 6, 83–96 (2006).

11. McKearnan SB, Wolfson J, Vock DM, Vazquez-Benitez G, O’Connor PJ. Performance of the Net Reclassification Improvement for Nonnested Models and a Novel Percentile-Based Alternative. Am J Epidemiol 187, 1327–1335 (2018).

12. Pencina MJ, D’Agostino RB, Massaro JM. Understanding increments in model performance metrics. Lifetime Data Anal 19, 202–218 (2013).

13. Usher-Smith JA, Sharp SJ, Luben R, Griffin SJ. Development and Validation of Lifestyle-Based Models to Predict Incidence of the Most Common Potentially Preventable Cancers. Cancer Epidemiol Biomarkers Prev 28, 67–75 (2019).

14. Mucci LA, et al. Familial Risk and Heritability of Cancer Among Twins in Nordic Countries. JAMA 315, 68–76 (2016).

15. Rashkin SR, et al. Pan-cancer study detects genetic risk variants and shared genetic basis in two large cohorts. Nat Commun 11, 4423 (2020).

16. Graff RE, et al. Cross-Cancer Evaluation of Polygenic Risk Scores for 17 Cancer Types in Two Large Cohorts. bioRxiv, Preprint at: https://doi.org/10.1101/2020.01.18.911578 (2020).

17. Shi Z, et al. Systematic evaluation of cancer-specific genetic risk score for 11 types of cancer in The Cancer Genome Atlas and Electronic Medical Records and Genomics cohorts. Cancer Med 8, 3196–3205 (2019).

18. Fritsche LG, et al. Association of Polygenic Risk Scores for Multiple Cancers in a Phenome-wide Study: Results from The Michigan Genomics Initiative. Am J Hum Genet 102, 1048–1061 (2018).

19. Amin Al Olama A, et al. Risk Analysis of Prostate Cancer in PRACTICAL, a Multinational Consortium, Using 25 Known Prostate Cancer Susceptibility Loci. Cancer Epidemiol Biomarkers Prev 24, 1121–1129 (2015).

20. Hoffmann TJ, et al. A large multiethnic genome-wide association study of prostate cancer identifies novel risk variants and substantial ethnic differences. Cancer Discov 5, 878–891 (2015).

21. Smith T, Gunter MJ, Tzoulaki I, Muller DC. The added value of genetic information in colorectal cancer risk prediction models: development and evaluation in the UK Biobank prospective cohort study. Br J Cancer 119, 1036–1039 (2018).

22. Garcia-Closas M, et al. Common genetic polymorphisms modify the effect of smoking on absolute risk of bladder cancer. Cancer Res 73, 2211–2220 (2013).

23. Mavaddat N, et al. Polygenic Risk Scores for Prediction of Breast Cancer and Breast Cancer Subtypes. Am J Hum Genet 104, 21–34 (2019).

24. Yang X, et al. Evaluation of polygenic risk scores for ovarian cancer risk prediction in a prospective cohort study. J Med Genet 55, 546–554 (2018).

25. Klein AP, et al. Genome-wide meta-analysis identifies five new susceptibility loci for pancreatic cancer. Nat Commun 9, 556 (2018).

26. Fritsche LG, et al. Exploring various polygenic risk scores for skin cancer in the phenomes of the Michigan genomics initiative and the UK Biobank with a visual catalog: PRSWeb. PLoS Genet 15, e1008202 (2019).

27. Abraham G, et al. Genomic risk score offers predictive performance comparable to clinical risk factors for ischaemic stroke. Nat Commun 10, 5819 (2019).

28. Khera AV, et al. Genetic Risk, Adherence to a Healthy Lifestyle, and Coronary Disease. N Engl J Med 375, 2349–2358 (2016).

29. Licher S, et al. Genetic predisposition, modifiable-risk-factor profile and long-term dementia risk in the general population. Nat Med 25, 1364–1369 (2019).

30. Brown KF, et al. The fraction of cancer attributable to modifiable risk factors in England, Wales, Scotland, Northern Ireland, and the United Kingdom in 2015. Br J Cancer 118, 1130–1141 (2018).

31. Brenner DR, et al. The burden of cancer attributable to modifiable risk factors in Canada: Methods overview. Prev Med 122, 3–8 (2019).

32. Poirier AE, et al. The current and future burden of cancer attributable to modifiable risk factors in Canada: Summary of results. Prev Med 122, 140–147 (2019).

33. Fry A, et al. Comparison of Sociodemographic and Health-Related Characteristics of UK Biobank Participants With Those of the General Population. Am J Epidemiol 186, 1026–1034 (2017).

34. National Lung Screening Trial Research T, et al. Reduced lung-cancer mortality with low-dose computed tomographic screening. N Engl J Med 365, 395–409 (2011).

35. De Koning H, Van Der Aalst C, Ten Haaf K, Oudkerk M. PL02.05 Effects of Volume CT Lung Cancer Screening: Mortality Results of the NELSON Randomised-Controlled Population Based Trial. Journal of Thoracic Oncology 13, S185 (2018).

36. Shieh Y, et al. Breast Cancer Screening in the Precision Medicine Era: Risk-Based Screening in a Population-Based Trial. J Natl Cancer Inst 109, (2017).

37. Hollands GJ, et al. The impact of communicating genetic risks of disease on risk-reducing health behaviour: systematic review with meta-analysis. BMJ 352, i1102 (2016).

38. Manichaikul A, Mychaleckyj JC, Rich SS, Daly K, Sale M, Chen WM. Robust relationship inference in genome-wide association studies. Bioinformatics 26, 2867–2873 (2010).

39. Huyghe JR, et al. Discovery of common and rare genetic risk variants for colorectal cancer. Nat Genet 51, 76–87 (2019).

40. Michailidou K, et al. Association analysis identifies 65 new breast cancer risk loci. Nature 551, 92–94 (2017).

41. Wooster R, Weber BL. Breast and ovarian cancer. N Engl J Med 348, 2339–2347 (2003).

42. Kazerouni N, Greene MH, Lacey JV, Jr., Mink PJ, Schairer C. Family history of breast cancer as a risk factor for ovarian cancer in a prospective study. Cancer 107, 1075–1083 (2006).

43. Olson SH, Kurtz RC. Epidemiology of pancreatic cancer and the role of family history. J Surg Oncol 107, 1–7 (2013).

44. Molina-Montes E, et al. Risk of pancreatic cancer associated with family history of cancer and other medical conditions by accounting for smoking among relatives. Int J Epidemiol 47, 473–483 (2018).

45. Schuz J, et al. European Code against Cancer 4th Edition: 12 ways to reduce your cancer risk. Cancer Epidemiol 39 Suppl 1, S1–10 (2015).

46. Demler OV, Paynter NP, Cook NR. Tests of calibration and goodness-of-fit in the survival setting. Stat Med 34, 1659–1680 (2015).

47. Gerds TA, Andersen PK, Kattan MW. Calibration plots for risk prediction models in the presence of competing risks. Stat Med 33, 3191–3203 (2014).

48. Benichou J, Gail MH. Estimates of absolute cause-specific risk in cohort studies. Biometrics 46, 813–826 (1990).

49. Ozenne B, Lyngholm Sørensen A, Scheike T, Torp-Pedersen C, Gerds TA. riskRegression: Predicting the Risk of an Event using Cox Regression Models. The R Journal 9, 440–460 (2017).

50. Sjolander A, Vansteelandt S. Doubly robust estimation of attributable fractions in survival analysis. Stat Methods Med Res 26, 948–969 (2017).

51. Dahlqwist E, Zetterqvist J, Pawitan Y, Sjolander A. Model-based estimation of the attributable fraction for cross-sectional, case-control and cohort studies using the R package AF. Eur J Epidemiol 31, 575–582 (2016).

