## Supplementary Tables and Figures for "Pan-cancer analysis demonstrates that integrating polygenic risk scores with modifiable risk factors improves risk prediction"

**Supplementary Table 1:** Characteristics of the UK Biobank study population, restricted to participants of predominately European ancestry, stratified by incident cancer status.

| Baseline Characteristic | Status at the end of follow-up | | | |
|---|---|---|---|---|
| | Cancer-free (n=390,998) | | Incident Cancer (n=22,755) | |
| | N | (%) | N | (%) |
| Age at assessment (years) | | | | |
| Mean (SD) | 56.53 | (8.02) | 60.19 | (6.88) |
| Sex | | | | |
| Females | 211513 | (54.10) | 11080 | (48.69) |
| Males | 179485 | (45.90) | 11675 | (51.31) |
| Smoking status | | | | |
| Never | 211936 | (54.20) | 10610 | (46.63) |
| Ever | 177685 | (45.44) | 12049 | (52.95) |
| Former | 137493 | (35.16) | 9174 | (40.32) |
| Current | 40192 | (10.28) | 2875 | (12.63) |
| Pack-years: Mean (SD) | 23.17 | (15.61) | 27.70 | (19.19) |
| Unknown | 1377 | (0.35) | 96 | (0.42) |
| Body-mass index (kg/m$^2$) | | | | |
| Normal: 18.5 ≤BMI<25 | 127817 | (32.69) | 6614 | (29.07) |
| Underweight: <18.5 | 1984 | (0.51) | 96 | (0.42) |
| Overweight: 25 ≤BMI<30 | 165859 | (42.42) | 10055 | (44.19) |
| Obese: BMI≥30 | 94070 | (24.06) | 5914 | (25.99) |
| Unknown | 1268 | (0.32) | 76 | (0.33) |
| Frequency of alcohol consumption | | | | |
| Never | 25864 | (6.61) | 1651 | (7.26) |
| Special occasions | 41557 | (10.63) | 2495 | (10.96) |
| 1-3 times per month | 43627 | (11.16) | 2312 | (10.16) |
| 1-2 times per week | 102999 | (26.34) | 5753 | (25.28) |
| 3-4 times per week | 94177 | (24.09) | 5225 | (22.96) |
| Daily | 82482 | (21.10) | 5304 | (23.31) |
| Unknown | 292 | (0.07) | 1651 | (7.26) |
| Ever diagnosed with hypertension? | | | | |
| Yes | 92541 | (23.67) | 6772 | (29.76) |
| Any first-degree relative diagnosed with cancer[1]? | | | | |
| Yes | 139833 | (35.76) | 9195 | (40.41) |
| Ever had screening for breast, prostate, or colorectal cancer? | | | | |
| Yes | 258249 | (66.05) | 16908 | (74.30) |
| Deaths occurring during follow-up | | | | |
| Death from any cause | 5982 | (1.53) | 4878 | (21.44) |
| Death due to cancer | 2103 | (0.54) | 4696 | (20.64) |

1. Based on self-reported cancers of the breast, prostate, lung, or bowel in non-adopted parents and siblings

**Supplementary Table 2**: Risk factors in addition to age and sex (if applicable), such as environmental exposures, lifestyle factors, and family history, that were included in the most comprehensive model for each cancer. Risk factors were selected based on literature review and availability in the UK Biobank cohort.

| Cancer Site | Risk Factors | Model Specification Notes |
|---|---|---|
| Prostate | Family history of prostate cancer | |
| Testis | - | |
| Breast | Family history of breast cancer, parity (≥1 live birth vs. none), age at menarche (years), menopausal status (pre-menopausal vs. post-menopausal vs. unknown or hysterectomy), ever used hormone replacement therapy (HRT), duration of oral contraceptive use (never used (0) vs. <20 years vs. ≥20 years), body mass index (BMI), weekly alcohol intake (grams)[1] | Interactions:<br>(BMI)*(Menopausal status) |
| Endometrium | Family history of cancer, parity (≥1 live birth vs. none), age at menarche (years), menopausal status (pre-menopausal vs. post-menopausal vs. unknown or hysterectomy), ever used HRT, duration of oral contraceptive use (never used (0) vs. <20 years vs. ≥20 years), BMI | |
| Ovary | Family history of breast cancer, parity (≥1 live birth vs. none), menopausal status (pre-menopausal vs. post-menopausal vs. unknown or hysterectomy), ever used HRT, duration of oral contraceptive use (never used (0) vs. <20 years vs. ≥20 years), BMI | Interactions:<br>(BMI)*(Menopausal status) |
| Cervix | Parity (≥1 live birth vs. none), duration of oral contraceptive use (never used (0) vs. <20 years vs. ≥20 years), cigarette pack-years | |
| Colorectum | Family history of bowel cancer, waist to hip ratio (WHR), cigarette pack-years, frequency of processed meat intake (<1 per week vs. ≥1 per week), moderate and/or strenuous physical activity (days per week), weekly alcohol intake (grams) | |
| Melanoma | Frequency of UV protection use (always vs. most times vs. sometimes vs. never out in the sun vs. never), time spent outside in the summer (hours per day), ease of tanning (very easily vs. moderate vs. mild vs. mostly burn) | |
| Lung | Family history of lung cancer, cigarettes per day (0 for never smokers), years of smoking (0 for never smokers), smoking status (never vs. former vs. current), $PM_{2.5}$ level in 2010 (micro-g/$m^3$) | Interactions:<br>(Former smoker)* (cigarettes/day)<br>(Former smoker)*(years of smoking) |
| Never-smokers | Family history of lung cancer, $PM_{2.5}$ level in 2010 (micro-g/$m^3$) | |
| Smokers | Family history of lung cancer, cigarettes per day, years of smoking, smoking status (former vs. current), years since quitting smoking (0 for current smokers), $PM_{2.5}$ level in 2010 (micro-g/$m^3$) | Interactions:<br>(Smoking status)*(cigarettes/day)<br>(Smoking status)*(years smoking) |
| NHL | - | |
| Bladder | BMI, smoking status (never vs. former vs. current), cigarette pack-years | |
| Kidney | BMI, smoking status (never vs. former vs. current), cigarette pack-years, ever diagnosed with hypertension | |
| Pancreas | BMI, smoking status (never vs. former vs. current), cigarette pack-years, family history of cancer (prostate, breast, lung or bowel) | |
| Oral cavity/pharynx | Smoking status (never vs. former vs. current), cigarette pack-years, weekly alcohol intake (grams) | |
| Lymphocytic leukemia | - | |
| Thyroid | BMI categories (BMI <25 vs. 25 ≤BMI <30, BMI≥30) | |

1. Weekly alcohol intake was derived by summing up the total number of drinks per week across different types of alcoholic beverages (beer, wine, spirits) and converting to units of alcohol based on values from UK Composition of foods integrated dataset: https://www.gov.uk/government/publications/composition-of-foods-integrated-dataset-cofid

**Supplementary Table 3**: Hazard ratios (HR) and corresponding p-values for each cancer risk factor estimated using a cause-specific Cox regression model accounting for death as a competing risk.

| Cancer Site and Risk Factors | HR[1] | (95% CI) | P-value |
|---|---|---|---|
| Prostate | | | |
| Family history of prostate cancer | 1.84 | (1.69 - 2.00) | $9.1\times10^{-46}$ |
| Breast | | | |
| Family history of breast cancer | 1.56 | (1.44 - 1.69) | $3.0\times10^{-29}$ |
| Parity (≥1 live birth) | 0.91 | (0.84 - 0.98) | 0.010 |
| Age at menarche (per 1 year) | 0.99 | (0.97 - 1.00) | 0.14 |
| BMI (per 1-unit increase) | 0.99 | (0.98 - 1.00) | 0.11 |
| Menopausal status: pre-menopausal | 1.00 | | |
| Menopausal status: post-menopausal | 0.36 | (0.24 - 0.53) | $1.8\times10^{-7}$ |
| Menopausal status: unknown/hysterectomy | 0.32 | (0.18 - 0.55) | $5.0\times10^{-5}$ |
| Ever used hormone replacement therapy (HRT) | 1.09 | (1.02 - 1.17) | $7.0\times10^{-3}$ |
| Oral contraceptive use: 0 (never used) | 1.00 | | |
| Oral contraceptive use: <20 years | 1.00 | (0.93 - 1.08) | 0.94 |
| Oral contraceptive use: ≥20 years | 1.10 | (0.98 - 1.23) | 0.12 |
| Alcohol intake[2] (70 g/week) | 1.04 | (1.02 - 1.05) | $2.3\times10^{-5}$ |
| BMI * menopausal status (post-menopausal) | | | $2.0\times10^{-5}$ |
| BMI * menopausal status (unknown/hysterectomy) | | | $7.6\times10^{-4}$ |
| Endometrium | | | |
| Family history of cancer | 1.11 | (0.95 - 1.30) | 0.20 |
| Parity (≥1 live birth) | 0.64 | (0.53 - 0.77) | $3.3\times10^{-6}$ |
| Age at menarche (per 1-year increase) | 0.92 | (0.88 - 0.97) | $1.8\times10^{-3}$ |
| BMI (per 1-unit increase) | 1.09 | (1.08 - 1.10) | $1.6\times10^{-49}$ |
| Menopausal status: pre-menopausal | 1.00 | | |
| Menopausal status: post-menopausal | 1.01 | (0.73 - 1.39) | 0.97 |
| Menopausal status: unknown/hysterectomy | 0.02 | (0.00 - 0.08) | $6.1\times10^{-8}$ |
| Ever used HRT | 0.84 | (0.71 - 0.99) | 0.041 |
| Oral contraceptive use: 0 (never used) | 1.00 | | |
| Oral contraceptive use: <20 years | 0.83 | (0.70 - 1.00) | 0.051 |
| Oral contraceptive use: ≥20 years | 0.36 | (0.24 - 0.56) | $4.9\times10^{-6}$ |
| Ovary | | | |
| Family history of breast cancer | 1.30 | (1.00 - 1.70) | 0.051 |
| Parity (≥1 live birth) | 0.72 | (0.57 - 0.91) | $6.2\times10^{-3}$ |
| BMI (per 1-unit increase) | 1.04 | (1.00 - 1.08) | 0.036 |
| Menopausal status: post-menopausal | 3.19 | (0.88 - 11.56) | 0.08 |
| Menopausal status: unknown/hysterectomy | 1.07 | (0.13 - 8.52) | 0.95 |
| Ever used HRT | 0.97 | (0.79 - 1.19) | 0.79 |
| Duration of oral contraceptive use: <20 years | 0.82 | (0.66 - 1.03) | 0.09 |
| Duration of oral contraceptive use: ≥20 years | 0.57 | (0.37 - 0.88) | 0.012 |
| BMI * menopausal status (post-menopausal) | | | 0.023 |
| BMI * menopausal status (unknown/hysterectomy) | | | 0.45 |
| Cervix | | | |
| Parity: ≥1 live birth | 1.81 | (1.30 - 2.53) | $4.9\times10^{-4}$ |

| | | | |
|---|---|---|---|
| Oral contraceptive use: 0 (never used) | | | |
| Oral contraceptive use: <20 years | 0.84 | (0.58 - 1.21) | 0.34 |
| Oral contraceptive use: ≥20 years | 1.08 | (0.69 - 1.69) | 0.74 |
| Cigarette pack-years (per 10 pack-years) | 1.15 | (1.05 - 1.25) | $1.8\times10^{-3}$ |
| Colon/rectum | | | |
| Family history of bowel cancer | 1.26 | (1.14 - 1.40) | $1.2\times10^{-5}$ |
| Waist to hip ratio (per 10% increase) | 1.17 | (1.11 - 1.24) | $2.2\times10^{-8}$ |
| Cigarette pack-years (per 10 pack-years) | 1.04 | (1.02 - 1.06) | $2.1\times10^{-4}$ |
| Processed meat intake: never | 1.00 | | |
| Processed meat intake: < once a week | 0.99 | (0.98 - 1.00) | 0.20 |
| Processed meat intake: ≥ once a week | 1.08 | (0.92 - 1.28) | 0.34 |
| Physical activity: strenuous or moderate (days/week) | 1.15 | (0.98 - 1.35) | 0.09 |
| Alcohol intake[2] (70 g/week) | 1.04 | (1.03 - 1.05) | $5.9\times10^{-9}$ |
| Melanoma | | | |
| Apply UV protection: never | 1.00 | | |
| Apply UV protection: sometimes | 1.37 | (1.10 - 1.69) | $4.1\times10^{-3}$ |
| Apply UV protection: most times | 1.82 | (1.48 - 2.25) | $2.1\times10^{-8}$ |
| Apply UV protection: always | 1.68 | (1.34 - 2.09) | $5.3\times10^{-6}$ |
| Apply UV protection: never in the sun | 1.06 | (0.46 - 2.42) | 0.89 |
| Time outdoors in the summer (hours per day) | 1.03 | (1.01 - 1.05) | $7.3\times10^{-3}$ |
| Ease of tanning: get very tan | 1.00 | | |
| Ease of tanning: moderate | 1.33 | (1.16 - 1.53) | $6.4\times10^{-5}$ |
| Ease of tanning: mild | 1.60 | (1.37 - 1.86) | $1.9\times10^{-9}$ |
| Ease of tanning: mostly burn | 1.61 | (1.37 - 1.89) | $4.0\times10^{-9}$ |
| Lung | | | |
| Family history of lung cancer | 1.61 | (1.43 - 1.81) | $7.4\times10^{-15}$ |
| $PM_{2.5}$ in 2010 (per 1 micro-g/m$^3$) | 1.10 | (1.05 - 1.15) | $1.9\times10^{-5}$ |
| Cigarettes per day | 1.00 | (0.99 - 1.00) | 0.52 |
| Years of smoking | 1.07 | (1.06 - 1.09) | $9.0\times10^{-23}$ |
| Smoking status: never | 1.00 | | |
| Smoking status: former | 0.34 | (0.25 - 0.46) | $6.9\times10^{-13}$ |
| Smoking status: current | 0.84 | (0.44 - 1.61) | 0.60 |
| Smoking status (former) * cigarettes per day | | | $2.2\times10^{-8}$ |
| Smoking status (former) * years of smoking | | | 0.40 |
| Lung (Never smokers) | | | |
| Family history of lung cancer | 0.91 | (0.61 - 1.38) | 0.67 |
| $PM_{2.5}$ in 2010 (per 1 micro-g/m$^3$) | 0.93 | (0.81 - 1.07) | 0.32 |
| Lung (Current or former smokers) | | | |
| Family history of lung cancer | 1.71 | (1.51 - 1.94) | $3.8\times10^{-17}$ |
| $PM_{2.5}$ in 2010 (per 1 micro-g/m$^3$) | 1.12 | (1.07 - 1.18) | $1.2\times10^{-6}$ |
| Cigarettes per day | 1.02 | (1.02 - 1.03) | $1.2\times10^{-13}$ |
| Years of smoking | 1.07 | (1.05 - 1.10) | $1.4\times10^{-8}$ |
| Smoking status: former | 1.00 | | |
| Smoking status: current | 2.12 | (0.92 - 4.89) | 0.08 |
| Years since quitting (per 1 year) | 1.01 | (0.98 - 1.04) | 0.44 |
| Smoking status (current) * cigarettes per day | | | $2.4\times10^{-8}$ |

| | | | |
|---|---|---|---|
| Smoking status (current) * years of smoking | | | 0.51 |
| Bladder | | | |
| Cigarette pack-years (per 10 pack-years) | 1.07 | (1.03 - 1.11) | $3.4\times10^{-4}$ |
| Smoking status: never | 1.00 | | |
| Smoking status: former | 1.52 | (1.26 - 1.82) | $8.0\times10^{-6}$ |
| Smoking status: current | 2.07 | (1.62 - 2.64) | $5.7\times10^{-9}$ |
| BMI (per 1-unit increase) | 1.01 | (0.99 - 1.02) | 0.45 |
| Kidney | | | |
| BMI (per 1-unit increase) | 1.04 | (1.02 - 1.05) | $1.7\times10^{-6}$ |
| Smoking status: never | 1.00 | | |
| Smoking status: former | 1.07 | (0.87 - 1.32) | 0.52 |
| Smoking status: current | 1.36 | (1.02 - 1.83) | 0.039 |
| Cigarette pack-years (per 10 pack-years) | 1.07 | (1.02 - 1.12) | $7.9\times10^{-3}$ |
| Diagnosed with hypertension | 1.69 | (1.44 - 1.98) | $2.1\times10^{-10}$ |
| Pancreas | | | |
| Family history of cancer (prostate, breast, lung, bowel) | 1.40 | (1.17 - 1.67) | $1.9\times10^{-4}$ |
| BMI (per 1-unit increase) | 1.03 | (1.01 - 1.05) | $8.4\times10^{-4}$ |
| Cigarette pack-years (per 10 pack-years) | 1.04 | (0.97 - 1.10) | 0.27 |
| Smoking status: never | 1.00 | | |
| Smoking status: former | 1.08 | (0.84 - 1.39) | 0.56 |
| Smoking status: current | 2.04 | (1.46 - 2.84) | $2.8\times10^{-5}$ |
| Oral cavity/pharynx | | | |
| Alcohol intake[2] (70 g/week) | 1.05 | (1.04 - 1.07) | $3.0\times10^{-10}$ |
| Cigarettes per day | 1.01 | (1.00 - 1.02) | $8.2\times10^{-3}$ |
| Years of smoking | 1.02 | (1.01 - 1.04) | $2.0\times10^{-4}$ |
| Smoking status: never | 1.00 | | |
| Smoking status: former | 0.58 | (0.37 - 0.91) | 0.050 |
| Smoking status: current | 1.09 | (0.60 - 1.96) | 0.43 |
| Thyroid | | | |
| BMI: <25 | 1.00 | | |
| BMI: 25 to <30 | 1.43 | (1.01 - 2.02) | 0.045 |
| BMI: 30 to <35 | 1.59 | (1.05 - 2.41) | 0.028 |
| BMI: ≥35 | 1.15 | (0.61 - 2.15) | 0.67 |

1. In addition to the listed risk factors all Cox regression models were adjusted for age and sex (if applicable)
2. Weekly alcohol intake was derived by summing up the total number of drinks per week across different types of alcoholic beverages (beer, wine, spirits) and converting to units of alcohol based on values from UK Composition of foods integrated dataset: https://www.gov.uk/government/publications/composition-of-foods-integrated-dataset-cofid

**Supplementary Table 4**: Hazard ratios (HR) per one standard deviation (SD) increase in the standardized polygenic risk score (PRS) and corresponding p-values were estimated using cause-specific Cox proportional hazards models, accounting for mortality as a competing risk. Results comparing three types of weighting approaches for combining individual risk variants in the PRS are presented: standard weights based on log odds ratios ($PRS_{\beta}$), unweighted sum of risk alleles ($PRS_{unw}$), and inverse variance (IV) weights ($PRS_{IV}$).

| Cancer Site | Cases | PRS Description | | HR[1] | (95% CI) | P-value | C index[2] | (C SE) | AUC[3] |
|---|---|---|---|---|---|---|---|---|---|
| | | Variants | Weights | | | | | | |
| Prostate | 4740 | 161 | $PRS_{\beta}$ | 1.39 | (1.35-1.43) | $2.0\times10^{-105}$ | 0.738 | (0.004) | 0.740 |
| | | | $PRS_{unw}$ | 1.66 | (1.62-1.71) | $3.4\times10^{-266}$ | 0.759 | (0.004) | 0.761 |
| | | | $PRS_{IV}$ | 1.77 | (1.72-1.82) | $4.3\times10^{-336}$ | 0.768 | (0.004) | 0.769 |
| Testis | 52 | 52 | $PRS_{\beta}$ | 2.18 | (1.66-2.87) | $2.3\times10^{-8}$ | 0.749 | (0.034) | 0.783 |
| | | | $PRS_{unw}$ | 1.96 | (1.49-2.58) | $1.4\times10^{-6}$ | 0.745 | (0.035) | 0.769 |
| | | | $PRS_{IV}$ | 2.26 | (1.71-2.99) | $1.0\times10^{-8}$ | 0.766 | (0.033) | 0.787 |
| Breast | 4760 | 162 | $PRS_{\beta}$ | 1.52 | (1.47-1.56) | $3.2\times10^{-183}$ | 0.632 | (0.005) | 0.637 |
| | | | $PRS_{unw}$ | 1.42 | (1.38-1.46) | $2.1\times10^{-129}$ | 0.618 | (0.005) | 0.623 |
| | | | $PRS_{IV}$ | 1.52 | (1.47-1.56) | $1.2\times10^{-180}$ | 0.635 | (0.004) | 0.637 |
| Endometrium | 643 | 9 | $PRS_{\beta}$ | 1.19 | (1.10-1.29) | $1.1\times10^{-5}$ | 0.749 | (0.011) | 0.755 |
| | | | $PRS_{unw}$ | 1.18 | (1.09-1.28) | $2.4\times10^{-5}$ | 0.749 | (0.011) | 0.754 |
| | | | $PRS_{IV}$ | 1.18 | (1.09-1.27) | $3.5\times10^{-5}$ | 0.749 | (0.011) | 0.754 |
| Ovary | 445 | 36 | $PRS_{\beta}$ | 1.13 | (1.04-1.24) | $6.2\times10^{-3}$ | 0.655 | (0.015) | 0.656 |
| | | | $PRS_{unw}$ | 1.18 | (1.07-1.29) | $5.8\times10^{-4}$ | 0.652 | (0.016) | 0.658 |
| | | | $PRS_{IV}$ | 1.20 | (1.10-1.32) | $9.0\times10^{-5}$ | 0.654 | (0.015) | 0.660 |
| Cervix | 282 | 10 | $PRS_{\beta}$ | 1.22 | (1.09-1.37) | $7.3\times10^{-4}$ | 0.750 | (0.017) | 0.745 |
| | | | $PRS_{unw}$ | 1.20 | (1.07-1.35) | $1.5\times10^{-3}$ | 0.749 | (0.017) | 0.745 |
| | | | $PRS_{IV}$ | 1.21 | (1.07-1.35) | $1.5\times10^{-3}$ | 0.749 | (0.017) | 0.745 |
| Colon/rectum | 2725 | 103 | $PRS_{\beta}$ | 1.32 | (1.27-1.37) | $5.5\times10^{-50}$ | 0.704 | (0.006) | 0.704 |
| | | | $PRS_{unw}$ | 1.46 | (1.41-1.52) | $9.2\times10^{-87}$ | 0.714 | (0.006) | 0.714 |
| | | | $PRS_{IV}$ | 1.48 | (1.43-1.54) | $1.8\times10^{-94}$ | 0.716 | (0.006) | 0.716 |
| Melanoma | 1805 | 24 | $PRS_{\beta}$ | 1.43 | (1.36-1.49) | $5.7\times10^{-51}$ | 0.663 | (0.008) | 0.652 |
| | | | $PRS_{unw}$ | 1.43 | (1.36-1.49) | $1.2\times10^{-50}$ | 0.662 | (0.008) | 0.652 |
| | | | $PRS_{IV}$ | 1.44 | (1.37-1.50) | $2.4\times10^{-53}$ | 0.664 | (0.008) | 0.654 |
| Lung | 1541 | 109 | $PRS_{\beta}$ | 1.16 | (1.11-1.22) | $1.5\times10^{-9}$ | 0.849 | (0.006) | 0.846 |
| | | | $PRS_{unw}$ | 1.15 | (1.09-1.20) | $1.5\times10^{-8}$ | 0.849 | (0.006) | 0.846 |
| | | | $PRS_{IV}$ | 1.17 | (1.12-1.23) | $1.2\times10^{-10}$ | 0.849 | (0.006) | 0.846 |
| NHL | 970 | 19 | $PRS_{\beta}$ | 1.16 | (1.09-1.24) | $1.0\times10^{-6}$ | 0.676 | (0.010) | 0.677 |
| | | | $PRS_{unw}$ | 1.18 | (1.11-1.25) | $2.9\times10^{-7}$ | 0.675 | (0.010) | 0.678 |
| | | | $PRS_{IV}$ | 1.15 | (1.08-1.22) | $1.1\times10^{-5}$ | 0.674 | (0.010) | 0.677 |
| Bladder | 890 | 15 | $PRS_{\beta}$ | 1.28 | (1.20-1.37) | $2.1\times10^{-13}$ | 0.813 | (0.008) | 0.803 |
| | | | $PRS_{unw}$ | 1.30 | (1.21-1.39) | $7.6\times10^{-15}$ | 0.814 | (0.008) | 0.803 |
| | | | $PRS_{IV}$ | 1.30 | (1.22-1.39) | $1.5\times10^{-15}$ | 0.814 | (0.008) | 0.804 |
| Kidney | 612 | 19 | $PRS_{\beta}$ | 1.16 | (1.08-1.26) | $1.0\times10^{-4}$ | 0.724 | (0.011) | 0.722 |
| | | | $PRS_{unw}$ | 1.13 | (1.05-1.22) | $1.5\times10^{-3}$ | 0.723 | (0.011) | 0.721 |

| | | | | | | | | | |
|---|---|---|---|---|---|---|---|---|---|
| | | | $PRS_{IV}$ | 1.15 | (1.07-1.24) | $2.6\times10^{-4}$ | 0.723 | (0.011) | 0.722 |
| Pancreas | 493 | 22 | $PRS_{\beta}$ | 1.49 | (1.36-1.62) | $1.3\times10^{-18}$ | 0.742 | (0.012) | 0.745 |
| | | | $PRS_{unw}$ | 1.44 | (1.31-1.57) | $1.1\times10^{-15}$ | 0.738 | (0.012) | 0.741 |
| | | | $PRS_{IV}$ | 1.49 | (1.37-1.63) | $5.2\times10^{-19}$ | 0.743 | (0.012) | 0.745 |
| Oral cavity/ pharynx | 481 | 14 | $PRS_{\beta}$ | 1.11 | (1.01-1.21) | $2.3\times10^{-2}$ | 0.686 | (0.015) | 0.702 |
| | | | $PRS_{unw}$ | 1.11 | (1.02-1.22) | $1.9\times10^{-2}$ | 0.686 | (0.015) | 0.702 |
| | | | $PRS_{IV}$ | 1.12 | (1.02-1.23) | $1.3\times10^{-2}$ | 0.687 | (0.015) | 0.702 |
| Lymphocytic leukemia | 340 | 75 | $PRS_{\beta}$ | 1.45 | (1.31-1.61) | $8.0\times10^{-13}$ | 0.735 | (0.016) | 0.719 |
| | | | $PRS_{unw}$ | 1.67 | (1.51-1.86) | $1.2\times10^{-21}$ | 0.755 | (0.015) | 0.736 |
| | | | $PRS_{IV}$ | 1.70 | (1.53-1.88) | $6.3\times10^{-23}$ | 0.756 | (0.015) | 0.738 |
| Thyroid | 191 | 12 | $PRS_{\beta}$ | 1.57 | (1.36-1.82) | $5.7\times10^{-10}$ | 0.666 | (0.023) | 0.679 |
| | | | $PRS_{unw}$ | 1.55 | (1.34-1.78) | $1.9\times10^{-9}$ | 0.671 | (0.023) | 0.676 |
| | | | $PRS_{IV}$ | 1.75 | (1.53-2.01) | $1.9\times10^{-15}$ | 0.692 | (0.022) | 0.701 |

1. Hazard ratio estimates are adjusted for age at assessment (years), sex (if applicable), family history of cancer (for sites with available self-reported information), genotyping array, the first 15 genetic ancestry principal components, and any additional risk factors applicable to each cancer listed in Supplementary Table 1
2. Harrell's C-index was calculated as a weighted average between 1 and 5 years of follow-up
3. AUC values were estimated at 5 years of follow-up

**Supplementary Table 5**: Assessment of model discrimination for each cancer comparing different combinations of conventional risk factors and polygenic risk scores (PRS).

| Cancer Site | Cases | Model specification | C index[2] | (C SE) | AUC[3] | Pseudo $R^2$ |
|---|---|---|---|---|---|---|
| Prostate | 4740 | Age | 0.710 | (0.004) | 0.713 | 0.349 |
| | | Age + family history | 0.716 | (0.004) | 0.720 | 0.366 |
| | | Age + $PRS_{IV}$ (instead of family history) | 0.763 | (0.004) | 0.766 | 0.496 |
| | | Age + family history + $PRS_{IV}$ | 0.768 | (0.004) | 0.769 | 0.510 |
| Testis | 52 | Age | 0.627 | (0.045) | 0.658 | 0.184 |
| | | Age + $PRS_{IV}$ | 0.766 | (0.033) | 0.787 | 0.605 |
| Breast | 4760 | Age | 0.543 | (0.005) | 0.548 | 0.017 |
| | | Age + family history | 0.559 | (0.005) | 0.562 | 0.031 |
| | | Age + $PRS_{IV}$ (instead of family history) | 0.620 | (0.005) | 0.626 | 0.122 |
| | | Age + family history + other predictors | 0.572 | (0.005) | 0.573 | 0.043 |
| | | Age + family history + other predictors + $PRS_{IV}$ | 0.635 | (0.004) | 0.637 | 0.146 |
| Endometrium | 643 | Age | 0.631 | (0.012) | 0.631 | 0.127 |
| | | Age + family history | 0.629 | (0.012) | 0.632 | 0.129 |
| | | Age + family history + other predictors | 0.744 | (0.011) | 0.747 | 0.463 |
| | | Age + family history + other predictors + $PRS_{IV}$ | 0.749 | (0.011) | 0.754 | 0.486 |
| Ovary | 445 | Age | 0.607 | (0.016) | 0.620 | 0.106 |
| | | Age + family history | 0.611 | (0.016) | 0.622 | 0.111 |
| | | Age + family history + other predictors | 0.641 | (0.015) | 0.643 | 0.151 |
| | | Age + family history + other predictors + $PRS_{IV}$ | 0.654 | (0.015) | 0.660 | 0.193 |
| Cervix[4] | 282 | Age | 0.729 | (0.017) | 0.719 | 0.346 |
| | | Age + other predictors | 0.736 | (0.018) | 0.731 | 0.386 |
| | | Age + other predictors + $PRS_{IV}$ | 0.749 | (0.017) | 0.745 | 0.437 |
| Colon/rectum | 2725 | Age + sex | 0.678 | (0.006) | 0.680 | 0.235 |
| | | Age + sex + family history | 0.679 | (0.006) | 0.681 | 0.239 |
| | | Age + sex + PRS (instead of family history) | 0.708 | (0.006) | 0.708 | 0.319 |
| | | Age + sex + family history + other predictors | 0.686 | (0.006) | 0.688 | 0.258 |
| | | Age + sex + family history + other predictors + $PRS_{IV}$ | 0.716 | (0.006) | 0.716 | 0.345 |
| Melanoma | 1805 | Age + sex | 0.597 | (0.008) | 0.592 | 0.063 |
| | | Age + sex + other predictors | 0.622 | (0.008) | 0.616 | 0.100 |
| | | Age + sex + other predictors + $PRS_{IV}$ | 0.664 | (0.008) | 0.654 | 0.180 |
| Lung | 1541 | Age + sex | 0.706 | (0.007) | 0.704 | 0.307 |
| | | Age + sex + family history | 0.713 | (0.007) | 0.714 | 0.333 |
| | | Age + sex + PRS (instead of family history) | 0.711 | (0.007) | 0.710 | 0.322 |
| | | Age + sex + family history + other predictors | 0.846 | (0.006) | 0.843 | 0.789 |
| | | Age + sex + family history + other predictors + $PRS_{IV}$ | 0.849 | (0.006) | 0.846 | 0.799 |
| Never smokers | 207 | Age + sex + family history + other predictors | 0.709 | (0.020) | 0.712 | 0.320 |
| | | Age + sex + family history + other predictors + $PRS_{IV}$ | 0.723 | (0.020) | 0.723 | 0.354 |
| Smokers | 1334 | Age + sex + family history + other predictors | 0.805 | (0.007) | 0.804 | 0.641 |

| | | | | | | |
|---|---|---|---|---|---|---|
| | | Age + sex + family history + other predictors + $PRS_{IV}$ | 0.809 | (0.007) | 0.808 | 0.657 |
| NHL | 970 | Age + sex | 0.667 | (0.010) | 0.669 | 0.207 |
| | | Age + sex + $PRS_{IV}$ | 0.674 | (0.010) | 0.677 | 0.227 |
| Bladder | 890 | Age + sex | 0.792 | (0.008) | 0.784 | 0.548 |
| | | Age + sex + other predictors | 0.808 | (0.008) | 0.796 | 0.595 |
| | | Age + sex + other predictors + $PRS_{IV}$ | 0.814 | (0.008) | 0.804 | 0.628 |
| Kidney | 612 | Age + sex | 0.685 | (0.012) | 0.687 | 0.253 |
| | | Age + sex + other predictors | 0.716 | (0.011) | 0.713 | 0.338 |
| | | Age + sex + other predictors + $PRS_{IV}$ | 0.723 | (0.011) | 0.722 | 0.366 |
| Pancreas | 493 | Age + sex | 0.692 | (0.014) | 0.695 | 0.273 |
| | | Age + sex + family history + other predictors | 0.714 | (0.013) | 0.715 | 0.336 |
| | | Age + sex + family history + other predictors + $PRS_{IV}$ | 0.743 | (0.012) | 0.745 | 0.439 |
| Oral cavity / pharynx | 481 | Age + sex | 0.610 | (0.014) | 0.627 | 0.117 |
| | | Age + sex + other predictors | 0.681 | (0.015) | 0.693 | 0.332 |
| | | Age + sex + other predictors + $PRS_{IV}$ | 0.687 | (0.015) | 0.702 | 0.356 |
| Lymphocytic leukemia | 340 | Age + sex | 0.695 | (0.016) | 0.688 | 0.255 |
| | | Age + sex + $PRS_{IV}$ | 0.756 | (0.015) | 0.738 | 0.415 |
| Thyroid | 191 | Age + sex | 0.577 | (0.024) | 0.590 | 0.060 |
| | | Age + sex + other predictors | 0.592 | (0.024) | 0.604 | 0.079 |
| | | Age + sex + other predictors + $PRS_{IV}$ | 0.692 | (0.022) | 0.701 | 0.310 |

1. Harrell's C-index was calculated as a weighted average between 1 and 5 years of follow-up
2. AUC values were estimated at 5 years of follow-up
3. $R^2$ values correspond to the measure of explained variation by Royston P (2006) [Reference 10]
4. Incorporating time-varying PRS effects did not have an impact on predictive performance (AUC=0.745). As an additional sensitivity analysis we estimated the AUC using scaled Schoenfeld residuals, to account for non-proportionality in the PRS effects, which yielded AUC=0.772.

**Supplementary Table 6**: Percentile net reclassification improvement (NRI) index comparing the most comprehensive conventional risk factor model for each cancer with the model that incorporates the polygenic risk score (PRS) in addition to these risk factors. Event NRI ($NRI_e$) and non-event NRI ($NRI_{ne}$) quantify reclassification improvement in cases and event-free individuals, respectively. Bootstrapped confidence intervals were obtained based on 1000 replicates.

| Cancer Site | NRI[1] | (95% CI) | $NRI_e$[2] | (95% CI) | $NRI_{ne}$[3] | (95% CI) |
|---|---|---|---|---|---|---|
| Prostate | 0.444 | (0.420, 0.467) | 0.441 | (0.417, 0.463) | 0.003 | (-0.0002, 0.007) |
| Testis | 0.368 | (0.174, 0.538) | 0.385 | (0.192, 0.555) | -0.018 | (-0.021, -0.014) |
| Breast | 0.384 | (0.365, 0.403) | 0.370 | (0.351, 0.388) | 0.014 | (0.010, 0.018) |
| Endometrium | 0.058 | (0.019, 0.098) | 0.060 | (0.021, 0.099) | -0.002 | (-0.004, 0.0003) |
| Ovary | 0.173 | (0.097, 0.236) | 0.168 | (0.093, 0.233) | 0.005 | (0.001, 0.008) |
| Cervix | 0.073 | (-0.001, 0.152) | 0.075 | (0.001, 0.156) | -0.002 | (-0.005, 0.001) |
| Colon/rectum | 0.273 | (0.244, 0.301) | 0.271 | (0.242, 0.298) | 0.003 | (0.0003, 0.005) |
| Melanoma | 0.293 | (0.260, 0.323) | 0.285 | (0.250, 0.315) | 0.008 | (0.005, 0.011) |
| Lung | 0.044 | (0.020, 0.070) | 0.029 | (0.004, 0.055) | 0.015 | (0.013, 0.017) |
| NHL | 0.119 | (0.078, 0.161) | 0.116 | (0.074, 0.157) | 0.004 | (0.002, 0.006) |
| Bladder | 0.116 | (0.069, 0.164) | 0.116 | (0.069, 0.164) | 0.0003 | (-0.002, 0.002) |
| Kidney | 0.065 | (0.026, 0.103) | 0.069 | (0.030, 0.107) | -0.004 | (-0.006, -0.003) |
| Pancreas | 0.228 | (0.174, 0.284) | 0.228 | (0.173, 0.284) | 0.001 | (-0.002, 0.003) |
| Oral cavity / pharynx | 0.065 | (0.022, 0.107) | 0.069 | (0.026. 0.111) | -0.004 | (-0.006, -0.002) |
| Lymphocytic leukemia | 0.307 | (0.227, 0.388) | 0.312 | (0.232, 0.394) | -0.004 | (-0.007, -0.002) |
| Thyroid | 0.415 | (0.330, 0.507) | 0.409 | (0.324, 0.501) | 0.006 | (0.003, 0.008) |

1. $NRI = NRI_e + NRI_{ne}$
2. Difference in proportions of subjects with events (incident cancer) correctly reclassified to a higher-risk category minus those reclassified to a lower-risk category
3. Difference in proportions of subjects without events (cancer-free) correctly reclassified to a lower-risk category minus those reclassified to a higher-risk category

**Supplementary Table 7**: P-values for differences in mean absolute risk between strata based on genetic and other risk factor profiles depicted in Figures 3, 4, and 5. Low polygenic risk score (PRS) corresponds to ≤20$^{th}$ percentile, average PRS is defined as >20$^{th}$ to <80$^{th}$ percentile, and high PRS includes individuals in the ≥80$^{th}$ percentile of the normalized PRS distribution. Individuals below the median of the modifiable risk factor distribution were classified as having reduced risk, whereas those above the median had elevated risk. Family history was based on self-reported cancers in first-degree relatives. All p-values are based on a two-sample t-test calculated for differences in risk at age 60 except for pre-menopausal breast cancer (mean risk at age 50).

| Contrast | Cancer Site | | | | | | | |
|---|---|---|---|---|---|---|---|---|
| | Prostate | Breast | Colon/rectal | Lung | | | | |
| Low PRS / Family history vs. Low PRS / No family history | $6.5\times10^{-37}$ | $1.4\times10^{-62}$ | $9.7\times10^{-26}$ | $2.2\times10^{-11}$ | | | | |
| Low PRS / Family history vs. Average PRS / No family history | $4.5\times10^{-25}$ | $4.6\times10^{-32}$ | $3.9\times10^{-60}$ | $1.6\times10^{-7}$ | | | | |
| Average PRS / No family history vs. Average PRS / Family history | $4.1\times10^{-119}$ | $4.0\times10^{-148}$ | $4.7\times10^{-62}$ | $6.8\times10^{-26}$ | | | | |
| Average PRS / Family history vs. High PRS / No family history | $5.9\times10^{-66}$ | $1.0\times10^{-78}$ | $1.0\times10^{-194}$ | $1.5\times10^{-7}$ | | | | |
| High PRS / No family history vs. High PRS / Family history | $4.4\times10^{-31}$ | $8.7\times10^{-49}$ | $2.0\times10^{-21}$ | $4.3\times10^{-13}$ | | | | |
| Low PRS / Family history vs. High PRS / No family history | - | - | - | 0.031 | | | | |
| Low PRS / No family history vs. Average PRS / No family history | - | - | - | $4.6\times10^{-10}$ | | | | |
| Average PRS / No family history vs. High PRS / No family history | - | - | - | $3.8\times10^{-16}$ | | | | |
| | Breast: Pre-menopausal | Breast: Post-menopausal | Colon/rectal | Melanoma | Lung | Bladder | Kidney | Oral/pharynx |
| Low PRS / Reduced modifiable vs. Low PRS / Elevated modifiable | $7.9\times10^{-20}$ | $3.5\times10^{-69}$ | $<10^{-324}$ | $1.4\times10^{-269}$ | $4.1\times10^{-192}$ | $1.3\times10^{-199}$ | $1.2\times10^{-318}$ | $2.8\times10^{-209}$ |
| Low PRS / Elevated modifiable vs. Average PRS / Reduced modifiable | $9.1\times10^{-59}$ | $1.2\times10^{-288}$ | $1.8\times10^{-42}$ | $1.3\times10^{-165}$ | $1.7\times10^{-181}$ | $7.9\times10^{-78}$ | $1.2\times10^{-224}$ | $6.0\times10^{-173}$ |
| Average PRS / Reduced modifiable vs. Average PRS / Elevated modifiable | $1.5\times10^{-36}$ | $6.3\times10^{-165}$ | $<10^{-324}$ | $<10^{-324}$ | $<10^{-324}$ | $<10^{-324}$ | $<10^{-324}$ | $<10^{-324}$ |
| Average PRS / Elevated modifiable vs. High PRS / Reduced modifiable | $2.4\times10^{-77}$ | $1.8\times10^{-122}$ | $7.8\times10^{-49}$ | $3.5\times10^{-139}$ | $<10^{-324}$ | $<10^{-324}$ | $<10^{-324}$ | $<10^{-324}$ |
| High PRS / Reduced modifiable vs. High PRS / Elevated modifiable | $3.4\times10^{-14}$ | $1.7\times10^{-40}$ | $3.6\times10^{-301}$ | $3.3\times10^{-211}$ | $1.9\times10^{-181}$ | $2.2\times10^{-179}$ | $1.9\times10^{-323}$ | $2.3\times10^{-203}$ |
| Low PRS / Elevated modifiable vs. Average PRS / Elevated modifiable | - | - | - | - | $1.1\times10^{-13}$ | $5.3\times10^{-100}$ | $3.6\times10^{-56}$ | $1.1\times10^{-12}$ |
| Average PRS / Elevated modifiable vs. High PRS / Elevated modifiable | - | - | - | - | $1.6\times10^{-19}$ | $5.6\times10^{-77}$ | $1.7\times10^{-52}$ | $1.6\times10^{-13}$ |
| Low PRS / Elevated modifiable vs. High PRS / Reduced modifiable | - | - | - | - | $2.7\times10^{-161}$ | 0.99 | $3.6\times10^{-98}$ | $7.3\times10^{-132}$ |
| Low PRS / Reduced modifiable vs. Average PRS / Reduced modifiable | - | - | - | - | $7.7\times10^{-128}$ | $1.6\times10^{-133}$ | $4.8\times10^{-88}$ | $3.4\times10^{-88}$ |
| Average PRS / Reduced modifiable vs. High PRS / Reduced modifiable | - | - | - | - | $1.4\times10^{-208}$ | $3.1\times10^{-79}$ | $5.2\times10^{-67}$ | $2.5\times10^{-65}$ |

**Supplementary Table 8:** Assessment of interaction on the absolute risk scale between ordinal polygenic risk score (PRS) categories (average: 20th to <80th percentile; high: ≥80th percentile vs. low: ≤20th percentile) and family history of cancer (yes vs. none) or elevated modifiable risk factor profile (summary score >50th percentile vs. ≤50th percentile). Coefficients and p-values for interaction terms were estimated using linear regression models with the predicted absolute risk of cancer at age 60 (age 50 for pre-menopausal breast cancer) as the outcome.

| Cancer Site | Interaction Combination | Interaction | | Overall Interaction P-value[1] |
|---|---|---|---|---|
| | | Coefficient | P-value | |
| Prostate | PRS(average) * family history(yes) | $7.8\times10^{-3}$ | $7.0\times10^{-15}$ | $9.0\times10^{-128}$ |
| | PRS(high) * family history(yes) | 0.026 | $7.4\times10^{-104}$ | |
| Breast | PRS(average) * family history(yes) | $3.0\times10^{-3}$ | $2.9\times10^{-9}$ | $1.2\times10^{-98}$ |
| | PRS(high) * family history(yes) | 0.011 | $6.4\times10^{-80}$ | |
| Colorectal | PRS(average) * family history(yes) | $3.3\times10^{-4}$ | 0.030 | $8.7\times10^{-14}$ |
| | PRS(high) * family history(yes) | $1.2\times10^{-3}$ | $2.8\times10^{-12}$ | |
| Lung | PRS(average) * family history(yes) | $-5.4\times10^{-5}$ | 0.88 | $7.0\times10^{-4}$ |
| | PRS(high) * family history(yes) | $1.1\times10^{-3}$ | $5.9\times10^{-3}$ | |
| Breast (Pre-menopausal) | PRS(average) * modifiable(elevated) | $1.3\times10^{-3}$ | 0.021 | $4.9\times10^{-7}$ |
| | PRS(high) * modifiable(elevated) | $3.6\times10^{-3}$ | $1.7\times10^{-7}$ | |
| Breast (Post-menopausal) | PRS(average) * modifiable(elevated) | $1.7\times10^{-3}$ | $4.8\times10^{-6}$ | $6.9\times10^{-24}$ |
| | PRS(high) * modifiable(elevated) | $4.6\times10^{-3}$ | $6.5\times10^{-24}$ | |
| Endometrium | PRS(average) * modifiable(elevated) | $5.0\times10^{-4}$ | $4.0\times10^{-5}$ | $5.3\times10^{-16}$ |
| | PRS(high) * modifiable(elevated) | $1.2\times10^{-3}$ | $1.2\times10^{-16}$ | |
| Ovary | PRS(average) * modifiable(elevated) | $1.5\times10^{-4}$ | $5.9\times10^{-7}$ | $1.4\times10^{-19}$ |
| | PRS(high) * modifiable(elevated) | $3.5\times10^{-4}$ | $1.9\times10^{-20}$ | |
| Cervix | PRS(average) * modifiable(elevated) | $7.7\times10^{-5}$ | 0.072 | $9.1\times10^{-5}$ |
| | PRS(high) * modifiable(elevated) | $2.2\times10^{-4}$ | $2.6\times10^{-5}$ | |
| Colorectum | PRS(average) * modifiable(elevated) | $1.0\times10^{-3}$ | $3.0\times10^{-38}$ | $1.3\times10^{-208}$ |
| | PRS(high) * modifiable(elevated) | $2.9\times10^{-3}$ | $1.3\times10^{-194}$ | |
| Melanoma | PRS(average) * modifiable(elevated) | $5.1\times10^{-4}$ | $3.8\times10^{-28}$ | $3.3\times10^{-122}$ |
| | PRS(high) * modifiable(elevated) | $1.3\times10^{-3}$ | $3.9\times10^{-118}$ | |
| Lung | PRS(average) * modifiable(elevated) | $1.2\times10^{-3}$ | $3.6\times10^{-8}$ | $1.1\times10^{-37}$ |
| | PRS(high) * modifiable(elevated) | $3.3\times10^{-3}$ | $4.3\times10^{-37}$ | |
| Bladder | PRS(average) * modifiable(elevated) | $3.4\times10^{-4}$ | $3.3\times10^{-11}$ | $1.5\times10^{-50}$ |
| | PRS(high) * modifiable(elevated) | $9.3\times10^{-4}$ | $1.2\times10^{-49}$ | |
| Kidney | PRS(average) * modifiable(elevated) | $1.8\times10^{-4}$ | $9.7\times10^{-9}$ | $5.5\times10^{-29}$ |
| | PRS(high) * modifiable(elevated) | $4.2\times10^{-4}$ | $1.6\times10^{-29}$ | |
| Pancreas | PRS(average) * modifiable(elevated) | $1.6\times10^{-4}$ | $3.7\times10^{14}$ | $2.1\times10^{-91}$ |
| | PRS(high) * modifiable(elevated) | $5.2\times10^{-4}$ | $6.5\times10^{-85}$ | |
| Oral cavity/pharynx | PRS(average) * modifiable(elevated) | $1.1\times10^{-4}$ | $9.6\times10^{-4}$ | $5.2\times10^{-11}$ |
| | PRS(high) * modifiable(elevated) | $2.7\times10^{-4}$ | $1.1\times10^{-11}$ | |
| Thyroid | PRS(average) * modifiable(elevated) | $4.7\times10^{-5}$ | $2.1\times10^{-4}$ | $5.4\times10^{-30}$ |
| | PRS(high) * modifiable(elevated) | $1.7\times10^{-4}$ | $6.6\times10^{-28}$ | |

1. Calculated based on a chi-square test with 2 degrees of freedom

**Supplementary Table 9**: Population attributable fractions (PAF) were estimated at 5 years of follow-up time for the top 20% (≥80[th] percentile) of the modifiable risk factor and polygenic risk score (PRS) distributions, respectively, and family history of cancer at the relevant site. PAF estimates and corresponding p-values were derived from Cox proportional hazard regression models that were adjusted for age at enrollment, sex, family history of cancer (if available), genotyping array, and the top 15 genetic ancestry principal components.

| | High Genetic Risk | | | High Modifiable Risk | | | Family History | | |
|---|---|---|---|---|---|---|---|---|---|
| | PAF | (95% CI) | P-value | PAF | (95% CI) | P-value | PAF | (95% CI) | P-value |
| Prostate | 0.232 | (0.215 - 0.249) | $5.5\times10^{-158}$ | - | - | - | 0.055 | (0.045 - 0.065) | $1.4\times10^{-25}$ |
| Testis | 0.303 | (0.135 - 0.472) | $4.5\times10^{-4}$ | - | - | - | - | - | - |
| Breast | 0.168 | (0.151 - 0.184) | $4.9\times10^{-87}$ | 0.037 | (0.019 - 0.054) | $6.4\times10^{-5}$ | 0.051 | (0.040 - 0.063) | $3.3\times10^{-18}$ |
| Pre-menopausal | 0.173 | (0.136 - 0.210) | $3.1\times10^{-20}$ | 0.004 | (-0.010 - 0.018) | 0.54 | 0.044 | (0.021 - 0.068) | $2.5\times10^{-4}$ |
| Post-menopausal | 0.159 | (0.139 - 0.179) | $4.\times10^{-54}$ | 0.044 | (0.019 - 0.071) | $6.9\times10^{-4}$ | 0.053 | (0.039 - 0.068) | $2.5\times10^{-13}$ |
| Endometrium | 0.043 | (0.002 - 0.084) | 0.039 | 0.353 | (0.303 - 0.404) | $1.5\times10^{-43}$ | 0.042 | (-0.020 - 0.103) | 0.18 |
| Ovary[1] | 0.082 | (0.031 - 0.134) | $1.6\times10^{-3}$ | 0.100 | (0.038 - 0.161) | $1.4\times10^{-3}$ | 0.025 | (-0.012 - 0.063) | 0.19 |
| Cervix | 0.123 | (0.057 - 0.190) | $2.7\times10^{-4}$ | 0.065 | (-0.001 - 0.130) | 0.053 | - | - | - |
| Colon/rectum | 0.167 | (0.145 - 0.190) | $9.2\times10^{-50}$ | 0.111 | (0.085 - 0.136) | $1.6\times10^{-17}$ | 0.027 | (0.012 - 0.042) | $5.3\times10^{-4}$ |
| Melanoma | 0.139 | (0.112 - 0.166) | $1.3\times10^{-23}$ | 0.066 | (0.039 - 0.093) | $1.1\times10^{-6}$ | - | - | - |
| Lung | 0.040 | (0.013 - 0.066) | $3.1\times10^{-3}$ | 0.636 | (0.606 - 0.666) | $1.4\times10^{-376}$ | 0.089 | (0.065 - 0.114) | $1.0\times10^{-12}$ |
| Never smokers[2] | 0.077 | (0.002 - 0.151) | 0.045 | 0.049 | (-0.076 - 0.174) | 0.44 | -0.013 | (-0.066 - 0.039) | 0.62 |
| Smokers | 0.035 | (0.007 - 0.063) | 0.015 | 0.663 | (0.620 - 0.706) | $3.2\times10^{-200}$ | 0.105 | (0.078 - 0.132) | $2.8\times10^{-14}$ |
| NHL | 0.053 | (0.020 - 0.087) | $1.9\times10^{-3}$ | - | - | - | - | - | - |
| Bladder | 0.085 | (0.048 - 0.121) | $4.7\times10^{-6}$ | 0.189 | (0.140 - 0.237) | $4.1\times10^{-14}$ | - | - | - |
| Kidney | 0.046 | (0.005 - 0.087) | 0.026 | 0.210 | (0.160 - 0.260) | $2.4\times10^{-16}$ | - | - | - |
| Pancreas[3] | 0.133 | (0.082 - 0.184) | $2.9\times10^{-7}$ | 0.118 | (0.064 - 0.172) | $1.9\times10^{-5}$ | 0.134 | (0.063 - 0.205) | $2.3\times10^{-4}$ |
| Oral cavity / pharynx | 0.006 | (-0.038 - 0.051) | 0.78 | 0.310 | (0.253 - 0.368) | $4.0\times10^{-26}$ | - | - | - |
| Lymphocytic leukemia | 0.269 | (0.204 - 0.334) | $7.4\times10^{-16}$ | - | - | - | - | - | - |
| Thyroid | 0.268 | (0.180 - 0.355) | $1.7\times10^{-9}$ | 0.202 | (0.039 - 0.366) | 0.015 | - | - | - |

1. Family history variable refers to self-reported breast cancer in a first-degree relative
2. The only modifiable risk factor is air pollution, modeled here as a categorical variable corresponding to $PM_{2.5}$ levels above the median
3. Family history variable refers to self-reported breast, prostate, lung, or bowel cancer in a first-degree relative

**Supplementary Figure 1:** Calibration plots comparing predicted and observed event probabilities for each of the 16 cancers examined. The most comprehensive risk factor model available is plotted in black and the same model with the addition of the polygenic risk score (PRS) is overlaid in red. Goodness of fit p-values ($P_{GoF}$) are based on the Hosmer & Lemeshow test.

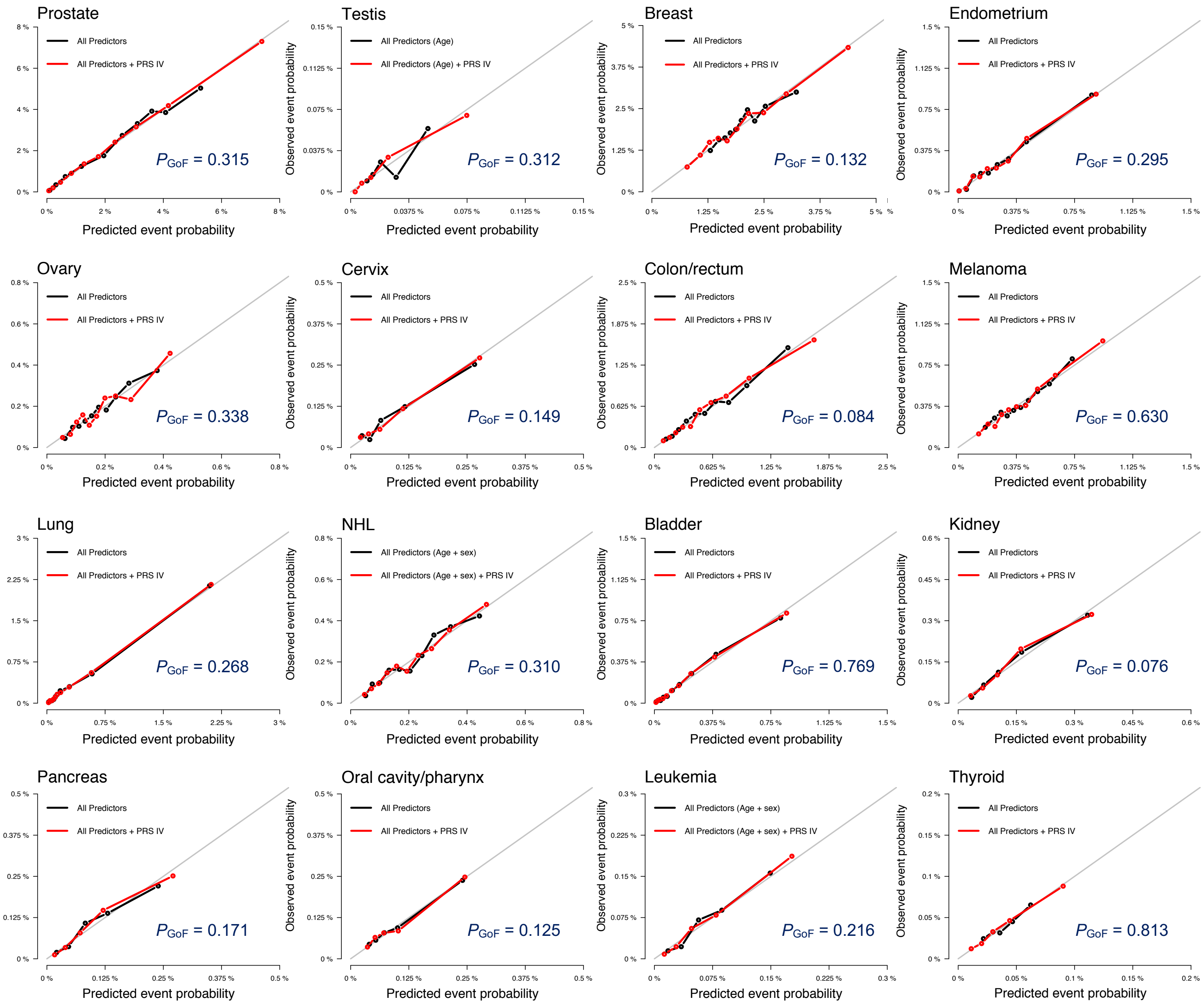

**Supplementary Figure 2:** Plots of Schoenfeld residuals and corresponding p-values for the cervical cancer polygenic risk score ($PRS_{IV}$). Below is a comparison of $PRS_{IV}$ effects estimated using a time-varying model (blue) and hazard ratio estimated under the proportionality assumption (red).

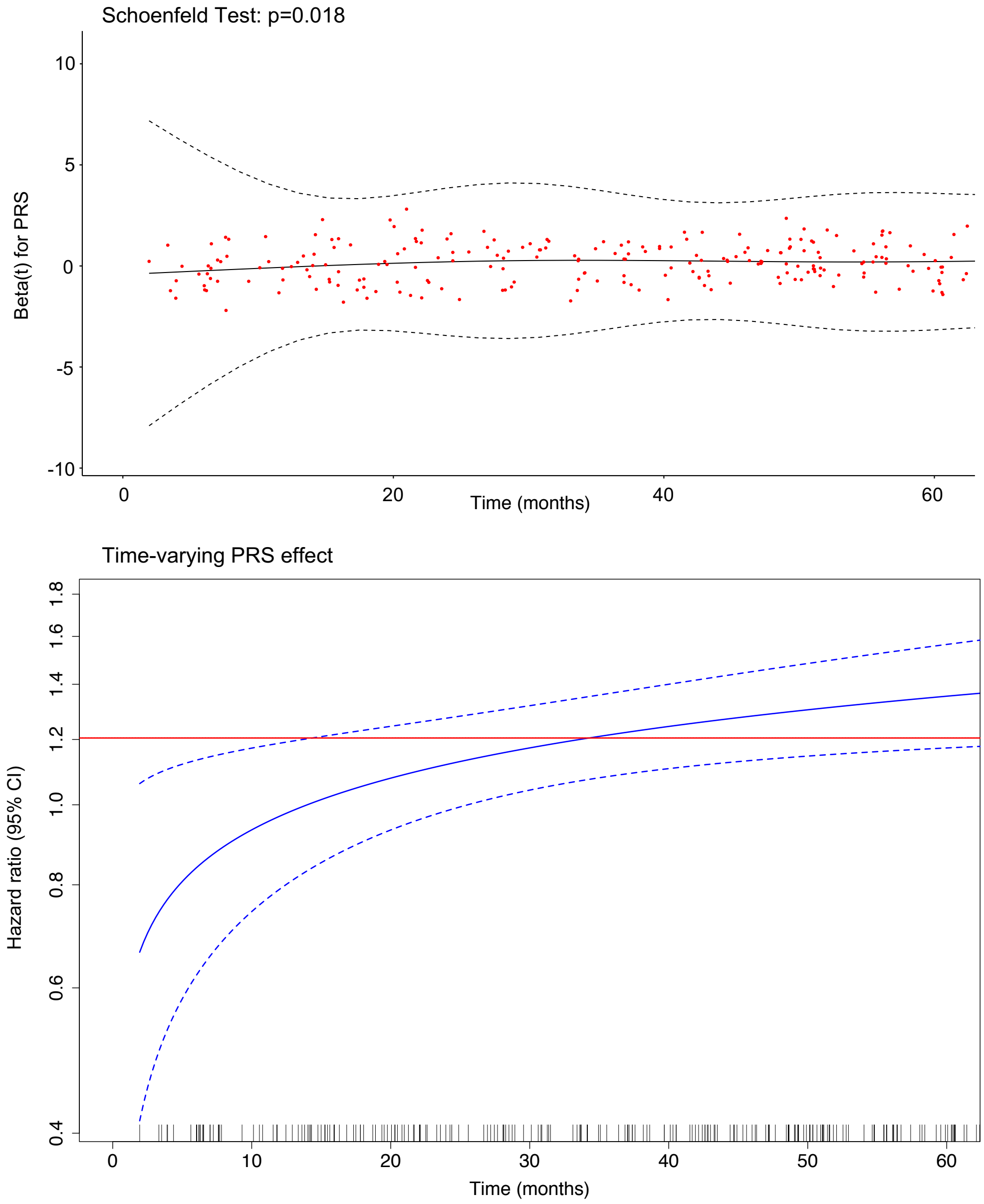

**Supplementary Figure 3:** Predicted 5-year absolute risks across strata defined by PRS and modifiable risk factors, where applicable, and age categories. Low genetic risk is based on percentiles of the standardized polygenic risk score (PRS). Low PRS corresponds to ≤20th percentile, average PRS is defined as >20th to <80th percentile, and high PRS includes individuals in the ≥80th percentile. Individuals below the median of the modifiable risk factor distribution were considered to have reduced risk, whereas those above the median had elevated risk. Absolute risks are visualized as box plots corresponding to the interquartile range (IQR), with outlying data points beyond 1.5*IQR plotted individually.

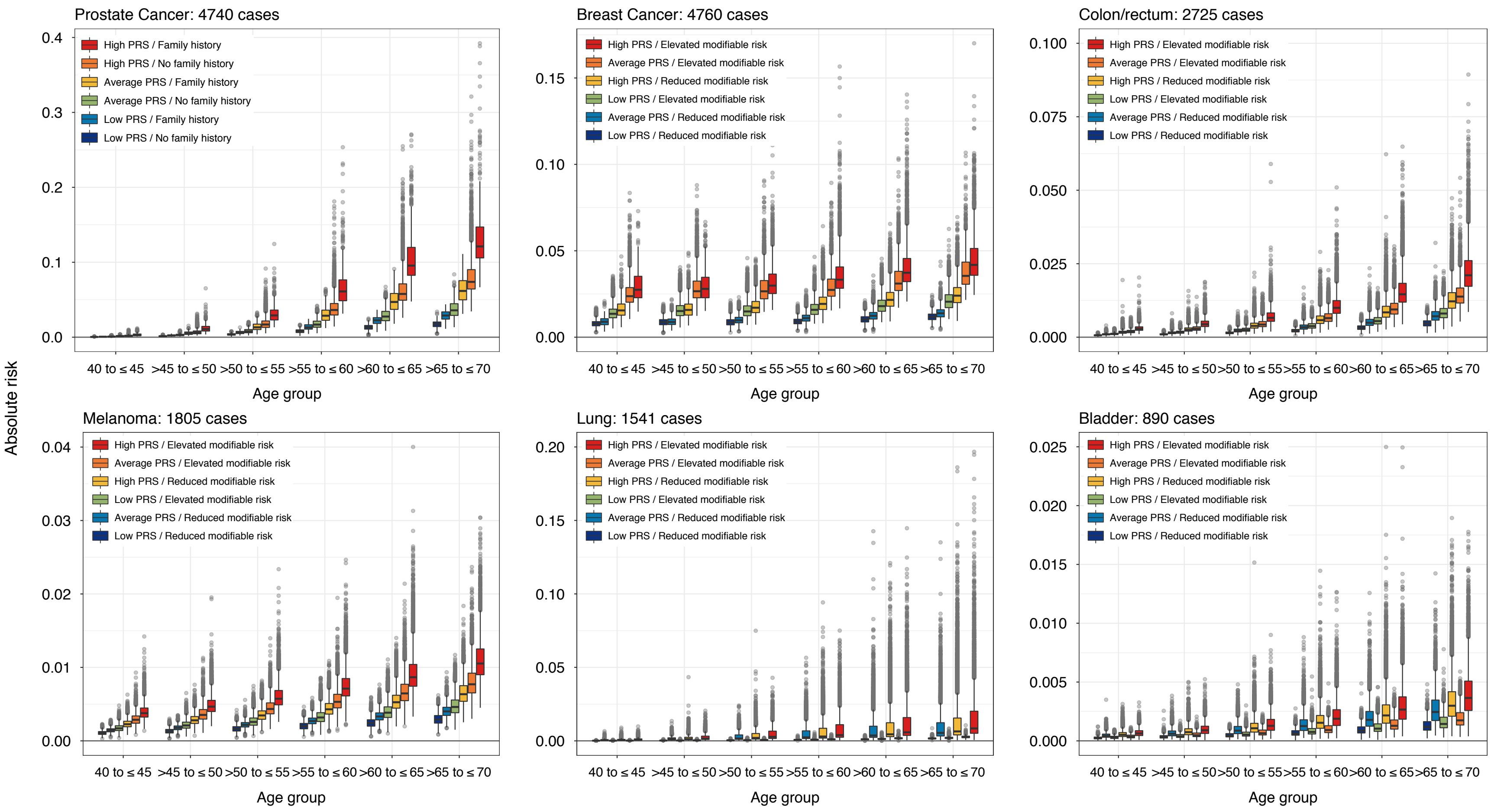

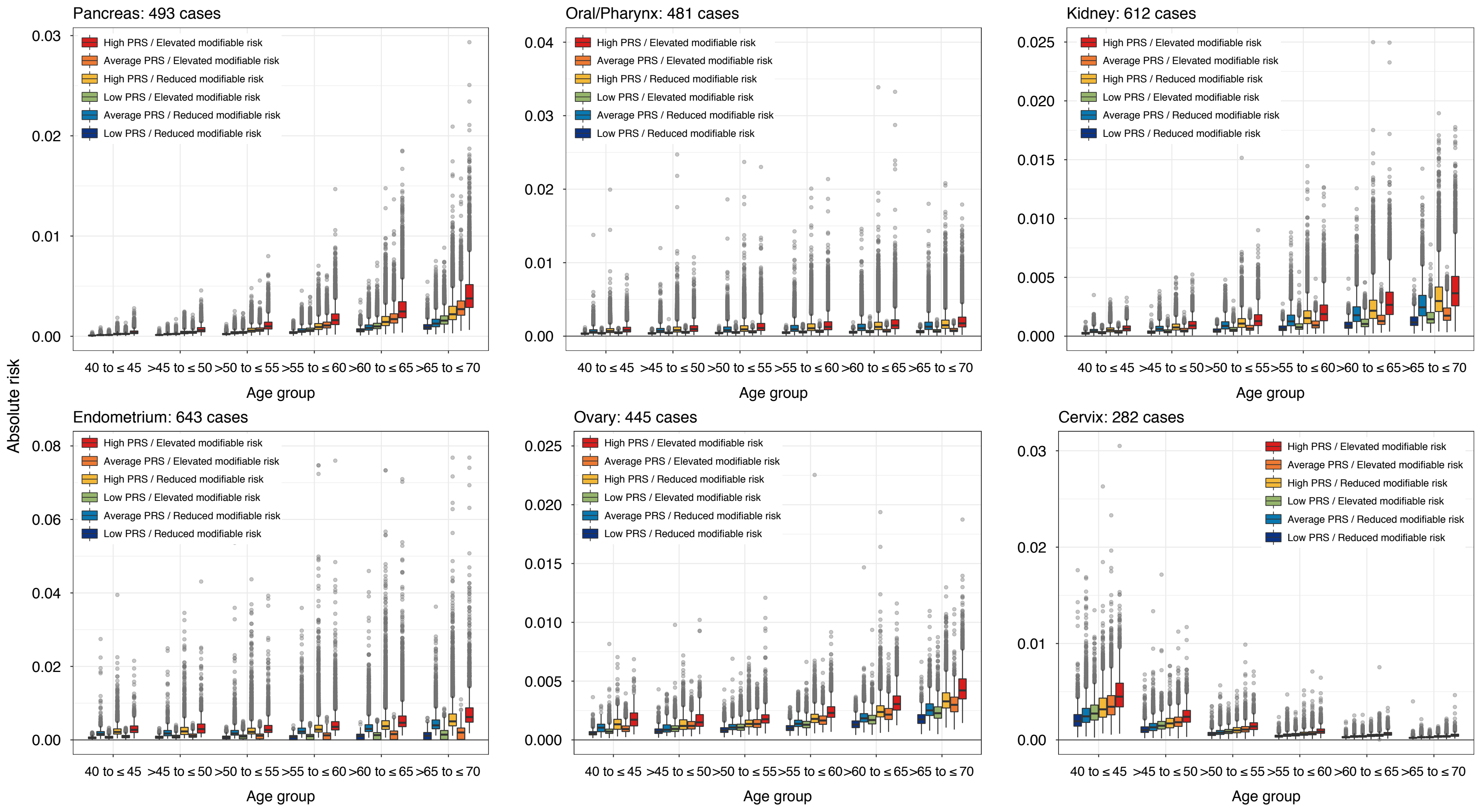
Pancreas: 493 cases
Oral/Pharynx: 481 cases
Kidney: 612 cases
Endometrium: 643 cases
Ovary: 445 cases
Cervix: 282 cases
High PRS / Elevated modifiable risk
Average PRS / Elevated modifiable risk
High PRS / Reduced modifiable risk
Low PRS / Elevated modifiable risk
Average PRS / Reduced modifiable risk
Low PRS / Reduced modifiable risk
Absolute risk
Age group
40 to ≤ 45
>45 to ≤ 50
>50 to ≤ 55
>55 to ≤ 60
>60 to ≤ 65
>65 to ≤ 70

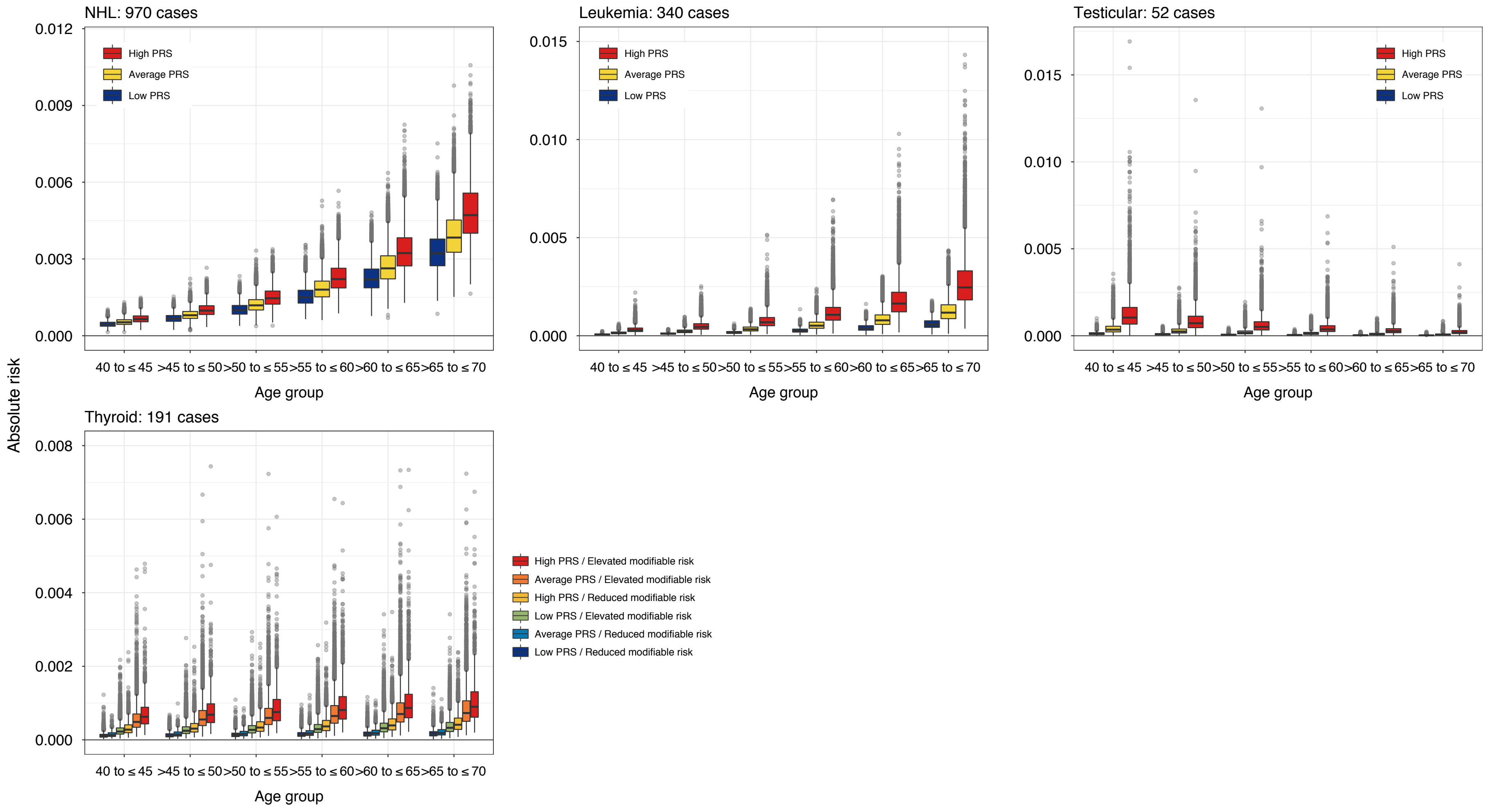
NHL: 970 cases
Leukemia: 340 cases
Testicular: 52 cases
Thyroid: 191 cases
Absolute risk
Age group
40 to ≤ 45
>45 to ≤ 50
>50 to ≤ 55
>55 to ≤ 60
>60 to ≤ 65
>65 to ≤ 70
High PRS
Average PRS
Low PRS
High PRS / Elevated modifiable risk
Average PRS / Elevated modifiable risk
High PRS / Reduced modifiable risk
Low PRS / Elevated modifiable risk
Average PRS / Reduced modifiable risk
Low PRS / Reduced modifiable risk
0.000
0.003
0.006
0.009
0.012
0.005
0.010
0.015
0.002
0.004
0.006
0.008

**Supplementary Figure 4:** Predicted 5-year absolute risk trajectories across strata defined by PRS and modifiable risk factors, where applicable. Low genetic risk is based on percentiles of the standardized polygenic risk score (PRS). Low PRS corresponds to ≤20th percentile, average PRS is defined as >20th to <80th percentile, and high PRS includes individuals in the ≥80th percentile. Individuals below the median of the modifiable risk factor distribution were considered to have reduced risk, whereas those above the median had elevated risk. P-values are based on t-tests comparing mean absolute risk in each stratum at age 60 or age 50 for cervical and testicular cancers. All statistical tests were two-sided.

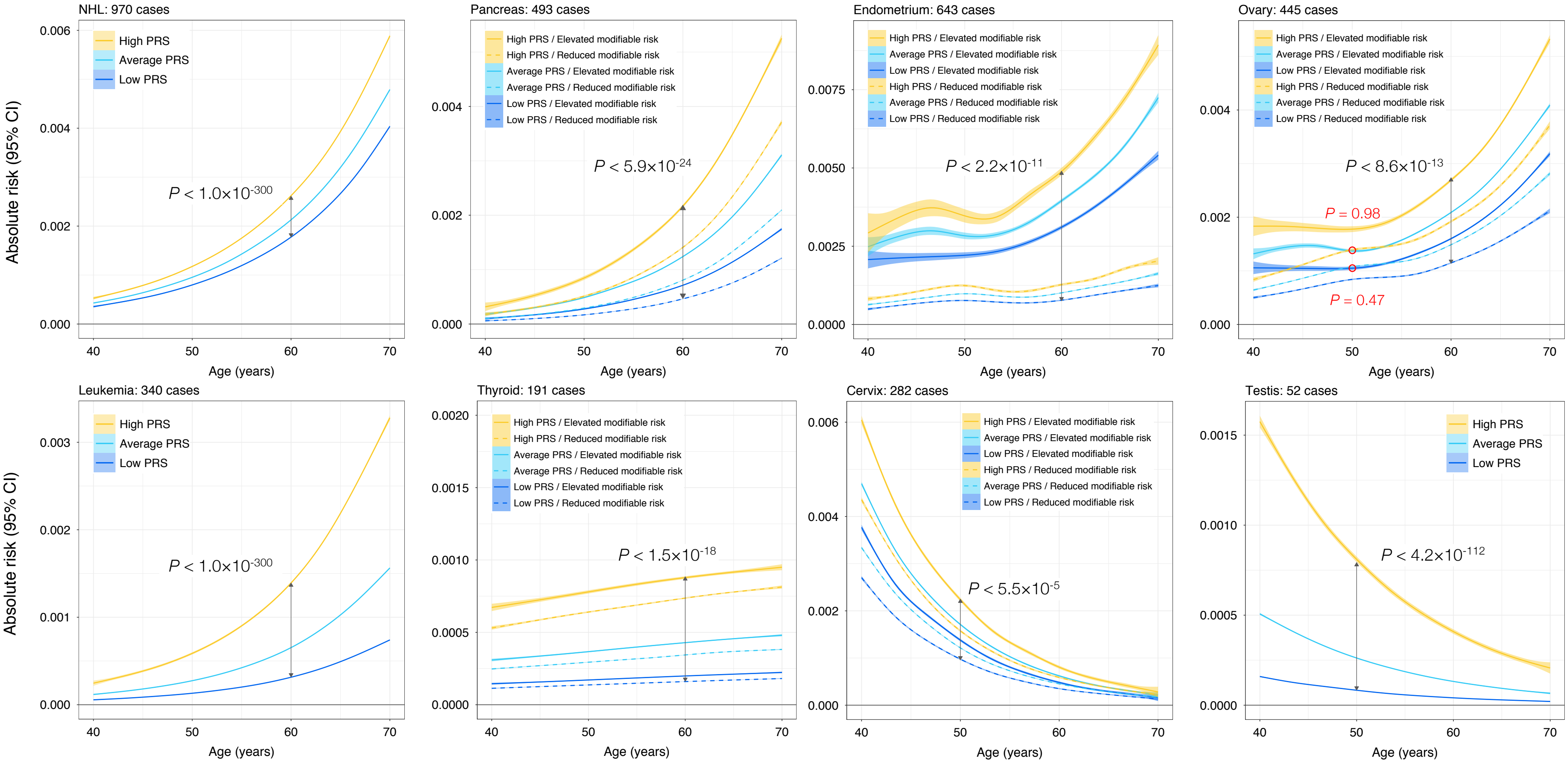


Note: for cervical cancer estimates of absolute risk were derived from the Cox proportional hazards model without time-varying PRS effects
